# Resting state fluctuations underlie free and creative verbal behaviors in the human brain

**DOI:** 10.1101/2020.03.01.971705

**Authors:** Rotem Broday-Dvir, Rafael Malach

## Abstract

Internally generated (free) ideas and creative thoughts constitute a fundamentally important aspect of the human experience, yet the neuronal mechanism driving these behaviors remains elusive. Here we examined the hypothesis that the common mechanism underlying free verbal behaviors is the ultra-slow activity fluctuations (termed “resting state fluctuations”) that emerge spontaneously in the human brain. In our experiment, participants were asked to perform three voluntary verbal tasks: a verbal fluency task, a verbal creativity task (alternative uses of everyday objects) and a divergent thinking task (instances of common concepts), during fMRI scanning. BOLD-activity during these tasks was contrasted with a control-deterministic verbal task, in which the behavior was fully determined by external stimuli. Our results reveal that in all three voluntary tasks, the verbal-generation responses displayed a gradual anticipatory buildup that preceded the deterministic control-related responses by ∼2 seconds. Importantly, variance analysis ruled out a time-jittered step-function response confound. Critically, the waveforms of the anticipatory buildups, as reflected in their time-frequency dynamics, were significantly correlated to the dynamics of resting state fluctuations, measured during a rest period prior to the tasks. Specifically, the amplitude of low frequency fluctuations (fALFF) of the resting state time-course and the voluntary verbal responses in the left inferior frontal gyrus (LH IFG), a central hub engaged in these tasks, were correlated across individual participants. This correlation was not a general BOLD-related or verbal-response related result, as it was not found during the externally-determined verbal control condition. Furthermore, it was specific to brain regions known to be involved in language production. These results indicate that the slow buildup preceding voluntary behaviors is linked to resting state fluctuations. Thus, these ubiquitous brain fluctuations may constitute a common neural mechanism underlying the generation of free and creative behaviors in the human brain.

## Introduction

Cognitive studies aiming at linking patterns of brain activity to behavior are typically characterized by a great effort of controlling the behavior as precisely as possible. However, such control overlooks a unique and important class of behaviors that are colloquially referred to as “free” or “voluntary” behaviors, i.e. behaviors that are internally generated, and are not directly driven by an external stimuli or instructions (Brass & Haggard, 2008; Haggard, 2008). Note that this definition is purely operational, and is unrelated to the deep and ongoing philosophical debate concerning free will. This operational distinction between deterministic and free behaviors is illustrated, for example, in studies that contrast voluntary self-initiated vs. cue-triggered or externally determined movements (Libet et al., 1983; Schurger et al., 2012). Another commonly used paradigm demonstrating this distinction is verbal fluency, in which participants perform freely-paced internal generation of verbal exemplars from a certain defined category (either semantic or phonemic, e.g. animals or words that start with the letter “a”). These blocks are compared to a deterministic control condition, e.g. wherein participants repeat a specific given word (Schlosser et al., 1998; Crosson et al., 2001; Knecht et al., 2003; Birn et al., 2010).

It may be argued that such category or domain-restricted behaviors are not completely free (e.g. involving a specific semantic category such as tools in a verbal fluency task). However, the critical difference between determined and free behaviors lies in the lack of a direct casual chain leading from an external cue to the specific behavior performed. Haggard (2008) defined three main components of this aspect: the “what” element, i.e. the instruction to repeat a specific word (“rest” for example) in the deterministic control condition, in contrast to freely recalling or selecting any specific tool from the entire category in a verbal fluency condition; the “when” element, i.e. repeating the word in response to an external cue in the deterministic condition, as compared to an internally-determined pace in free verbal fluency; and the “whether” element – the actual occurrence of a behavior directly stems from an external trigger in the control condition, versus free internally-driven generation in verbal fluency. This break in the external-causal chain which typifies the free condition raises the question of mechanism. For example, what neuronal process may induce the participants to select a specific tool from the entire category-i.e. to select a specific decision out of the numerous potential alternatives?

While such voluntary behaviors are of critical importance to the sense of spontaneity and freedom which are essential elements of human wellbeing, not less important is the realization that the phenomena of creative insights and original thinking all emerge, by definition, through a free and internally generated process, that can never be instructed deterministically from outside. Thus, insight into a neuronal mechanism underlying free behavior may bear a direct impact on our understanding of a fundamental cognitive ability underlying human progress in all scientific and cultural domains. The topic of the brain mechanisms linked to creativity has been of great interest, as well as controversy regarding how to probe this “impossible” quest of studying creativity inside the laboratory (Fink et al., 2007; Dietrich & Kanso, 2010; Jung et al., 2013; Benedek, Jauk, et al., 2014; Beaty et al., 2019). The brain networks involved in such behaviors, specifically in the commonly employed divergent thinking tasks, have been extensively mapped, demonstrating among other regions the consistent involvement of the lateral prefrontal cortex, the anterior cingulate cortex and the default mode network (DMN) (Fink et al., 2009; Abraham et al., 2012; Gonen-Yaacovi et al., 2013; Mayseless et al., 2015; Wu et al., 2015; Beaty et al., 2016). A few studies have examined more directly the brain activity preceding a creative idea-generation or problem solving “a-ha” moment, revealing different activation patterns during and prior to riddle solving by an “insight” experience as compared to non-insight (Jung-Beeman et al., 2004; Kounios et al., 2006).

However, the specific neural mechanisms and dynamics driving these moments of sudden “out-of-the-blue” creative, perhaps world-changing, ideas, are still an unresolved and intriguing question. Indeed, compared to the vast research effort that has been directed towards controlled, deterministic behaviors, far fewer studies attempted to decipher the neuronal mechanism underlying free behavior.

An important and pioneering line of research began with the seminal discovery of the “readiness potential” (Kornhuber & Deecke, 1965). This slow buildup of EEG signal prior to motor movements was shown to specifically precede voluntary movements rather than externally-determined ones. Elegant subsequent work by Libet et al. (1983), as well as more recent works (Soon et al., 2008; Fried et al., 2011; Schultze-Kraft et al., 2016) demonstrated that this buildup occurs mainly below the threshold of awareness, leading to a lively philosophical and psychological debate regarding free will (Frith et al., 2000; Wegner, 2017; Maoz et al., 2019). Most studies considered the readiness potential as reflecting a sub-conscious goal-directed deliberation process. However, an intriguing alternative has been recently put forward by Schurger et al. (2012), who hypothesized that the readiness potential actually corresponds to a stochastic fluctuation generated by accumulation of neuronal noise.

Interestingly, this gradual anticipatory signal is not solely a motor-related effect, and is also observed a few seconds before free recall of previously displayed visual stimuli, but not before their direct viewing (Polyn et al., 2005; Gelbard-Sagiv et al., 2008; Norman et al., 2019). Extending on Schurger et al. (2012), we have hypothesized (Moutard et al., 2015) that in fact the entire range of free behaviors, from motor decisions regarding when to act, to moments of creative insight, all depend on slow stochastic fluctuations as their driving mechanism.

It should be noted that if this hypothesis is correct, then these stochastic fluctuations must be an extremely ubiquitous brain phenomena, since free behaviors can be found in a widely diverse set of tasks ranging from motor movements, visual imagery, memory recall, creative verbal generation tasks and even music improvisation, to name a few (e.g. Libet et al., 1983; Finke, 1996; Limb & Braun, 2008; Benedek, Beaty, et al., 2014; Norman et al., 2019). A second, obvious requirement of such a neural mechanism is that it should be active spontaneously in the absence of external stimuli or deterministic instructions. Intriguingly, a ready neuronal candidate fulfilling these requirements is glaringly evident in the phenomena of ultra-slow spontaneous (also termed resting state) activity fluctuations. These slow (typically <0.1 Hz) intrinsic fluctuations were found across the entire human cortex, using a range of methods, from fMRI (Biswal et al., 1995; Nir et al., 2006; Fox & Raichle, 2007) to single neuron and IEEG recordings (Nir et al., 2008). Spontaneous fluctuations are of great interest, as they were shown to be linked to task related activation patterns and individual characteristics (Smith et al., 2009; Harmelech & Malach, 2013; Tavor et al., 2016; Grossman et al., 2019). Specifically, previous studies have shown that different resting state activity parameters of individual participants, including functional connectivity patterns, fractional amplitude of low frequency fluctuations (fALFF), entropy and regional homogenity measures are correlated with their creative behavior scores and abilities, insight solutions quantity, and free verbal-generation performance levels (Kounios et al., 2008; Takeuchi et al., 2012; Beaty et al., 2014; Wei et al., 2014; Yin et al., 2015; Beaty et al., 2018; Shi et al., 2019; Sun et al., 2019). However, despite this extensive research, the functional role that the resting state fluctuations may play in these free behaviors remains a mystery.

Here, we propose that resting state activity fluctuations drive the generation of verbal or creative ideas, by raising the neural activity in task-relevant networks above the decision bound and thus eliciting the free behavior events. This hypothesis is illustrated schematically in figure 1 A. Specifically, we hypothesize that resting state fluctuations can explore different possible verbal solutions by stochastically activating a diverse range of behaviorally relevant networks. When such stochastic activity crosses an activation threshold, this leads to the conscious emergence of a word or a verbal idea in the mind. Because of the inherently slow nature of resting state fluctuations, the stochastic accumulation leading to the threshold crossing will be appear as a slow, gradual build-up of activity preceding the event onset. A crucial prediction here is that if indeed a resting state fluctuation actually constitutes the anticipatory buildup, then there should be a tight correlation between the waveforms of resting state fluctuations, measured during rest, and the anticipatory buildup found prior to the free behavior. Thus, if there are substantial individual differences in the shape or dynamics of resting state fluctuations, then these individual differences should also be reflected in the buildup appearing prior to free behaviors. This prediction is schematically illustrated in figure 1B: individuals whose spontaneous fluctuations display very slow dynamics, will also show a very slow and sluggish anticipatory buildup prior to the voluntary event, as visualized in the left column time-courses in figure 1B. In contrast, individuals whose spontaneous fluctuations are faster and steeper, will also show a sharper buildup prior to voluntary behaviors (right column in figure 1B).

**Figure 1.**
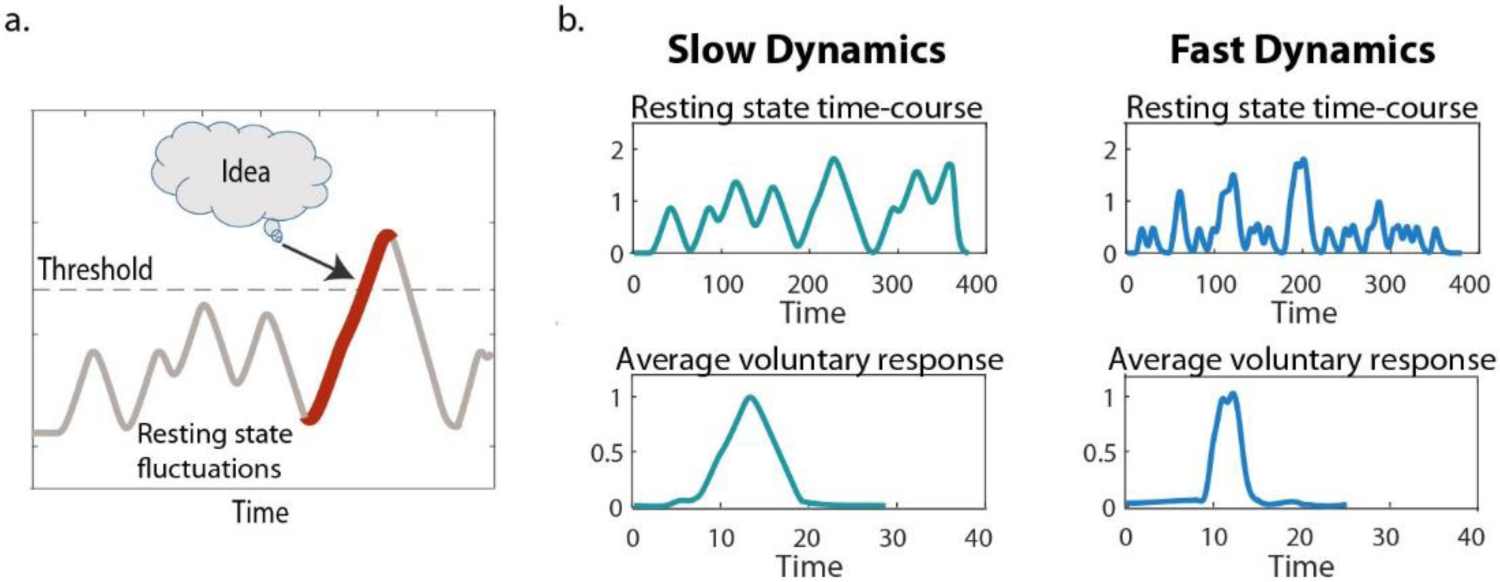
A. A scheme of the hypothesis: resting-state fluctuations drive the formation of a new verbal idea. This effect is manifested as a gradual signal buildup, leading to the crossing of a threshold, and by so bringing the idea into conscious thought. B. Schematic representation of BOLD resting state fluctuations and average event-related voluntary responses with slow dynamics (left column) and fast dynamics (right column).

We further hypothesize that free behaviors, including creative insights, are all similarly driven by resting state fluctuations. Hence, we predict to see a link between free behavior responses and resting state fluctuations across all tasks tested in the study, but, importantly, not between resting state fluctuations and externally determined behavior responses.

In the present study, we examined the hypothesis in the verbal-language domain, in a BOLD-fMRI experiment. Participants underwent a resting state scan, after which they completed three different free verbal tasks: verbal fluency, creativity and divergent thinking tasks, as well as an externally-driven verbal repetition task. By allowing long trial durations and extracting verbal events that are isolated in time, we were able to examine the neural activity that precedes the creative event onset, and compare it to the control, externally-determined events as well as to resting state fluctuations of individual participants. Our results reveal a significant and selective anticipatory buildup prior to all voluntary, but not externally determined, verbal behaviors. Furthermore, they demonstrate a significant correlation between free verbal and creative idea generation events and resting state activity dynamics, but not between resting state and the externally driven control events.

## Results

Our results were obtained in 23 participants in total. The participants first underwent an 8-minute resting state scan in which they were instructed to rest with open eyes, while maintaining their gaze in the center of a gray screen. This rest scan was then followed by three different voluntary verbal behavior tasks conducted inside the fMRI. We measured the BOLD signal, as well as pupil size and eye movements, while the participants completed a verbal fluency (VF) task (n=22), an alternative uses (AU) task (n=20) and a common instances (INST) task (n=15), performed in two separate scanning sessions. Each fMRI scanning session began with the resting state scan, followed by either 5 runs of the VF task, or 2 AU runs and 2 INST runs, in a counterbalanced order. Figure 2 depicts the three tasks as well as the control condition, which was the same in all experimental runs. In the VF condition, participants covertly generated exemplars from a specific phonemic category (words that begin with a certain Hebrew letter) or semantic category (e.g. tools, birds etc.). They reported each exemplar generation event by pressing a button, after which a short auditory cue was heard. Each VF block lasted 2.5 minutes and was terminated by the visual cue “Break”, followed by a 30-second break period. During the control, externally determined blocks, participants were instructed to wait until an auditory cue was played, and then to covertly repeat a certain word, e.g. “rest”, that was specified in the beginning of each control block, and immediately afterwards press the button. The auditory cues in the control blocks replayed the participants’ inter-response intervals during the VF blocks, resulting in equally spaced externally-generated (control) events and internally-generated (VF) events. Each VF run included one semantic and one phonemic verbal generation blocks, as well as two control blocks, in random order.

**Figure 2.**
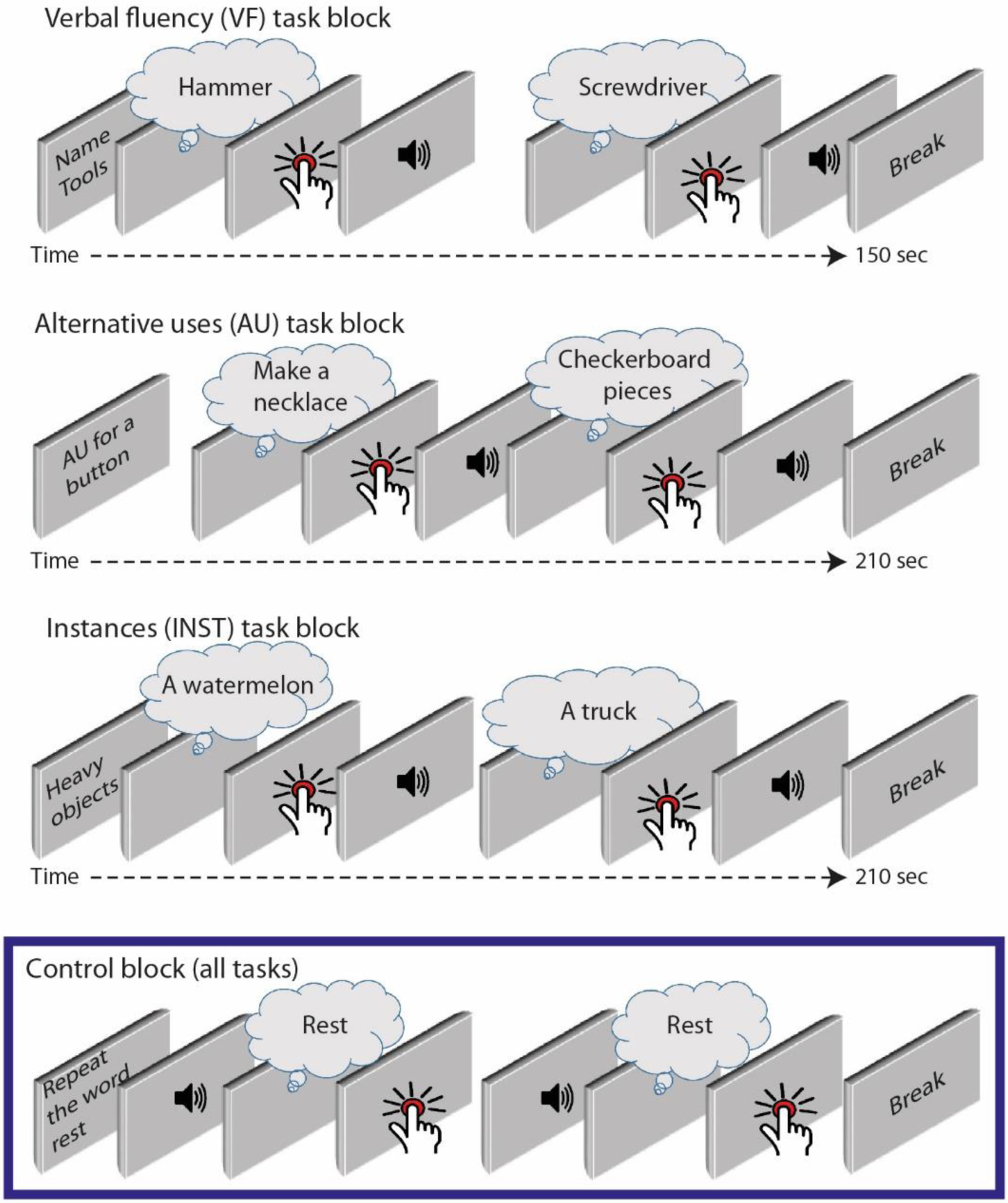
A scheme of the three experimental task blocks and the control block. VF blocks were initiated with a visual cue instructing the participant what category of words to generate. The participants then had to covertly generate words that belonged to the specific category, and press a button immediately each time an idea came to mind. They continued trying to think of relevant words until the block terminated after ∼150 seconds, with the cue “Break”, followed by a 30-second break. AU blocks were designed similarly to the VF blocks, but were longer and lasted ∼210 seconds. The participants’ task was to think of creative alternative uses to an everyday object that was specified at the beginning of each block. Instances task blocks also had the same structure as the two former tasks, with the instruction here being to generate as many instances of common concepts as possible until the block terminates. The control blocks were identical in all 3 experiments. They were initiated by the instruction to covertly repeat a specific word that was different in each block, after each time an auditory cue was heard. After the covert repetition of the word, the participants pressed the button. Control blocks were ∼120 seconds long, and the timings of the auditory cues replayed the participants’ performance in the 120 final seconds of the previous experimental block.

The AU task, a classic creative thinking task, required the participants to think of original and creative uses for everyday objects, like a button or a paperclip (Guilford, 1967; Kaufman et al., 2008). AU blocks had a similar structure to the VF blocks, in which the participants pressed a button each time a new idea came to their mind, after which an auditory cue was played. The final task, the INST task, had a similar structure to the two previous tasks, yet here the participants were instructed to generate diverse examples for common instances, including things that are either heavy, loud, round or tall (Silvia, 2011). Control blocks in the AU and INST experiments were identical to those in the VF experiment, requiring “passive” externally-cued word repetition, replaying the participants’ performance timings in the voluntary blocks. All three tasks included 2 experimental blocks and 2 control blocks in each run, and were performed in a covert manner, in order to avoid movement-related artifacts due to overt speech.

By setting long block durations, we were able to obtain creative and control events that were separated in time with a relatively long “no-event” incubation period preceding them, of at least 12-seconds. This allowed us to inspect the neural activity leading up to the subjective experience of a spontaneous emergence of an idea or a verbal exemplar, as compared to the activity preceding control passive word production in response to an external cue, in the absence of confound activations from a previous event.

### Behavioral results

A number of parameters of the participants’ behavior were examined, including their performance, quantified as the total number of words or creative ideas they produced, and the temporal dynamics of their production rate across the experimental blocks duration. Participants produced on average 20.64±5.0 exemplars in each verbal fluency block, 9.75±6.22 ideas in each alternative uses block, and 23.13±9.91 items in every instances block. One-way ANOVA analysis followed by post-hoc comparisons revealed that the number of words produced in the AU blocks was significantly lower than in the other two tasks, and that the performance in the VF and INST tasks was not statistically different (F(2,61) = 14.81, p<0.0001; Bonferroni post-hoc comparisons: VF>AU, p<0.001; INST>AU, p<0.001; VF∼= INST, p=0.875). Supplementary figure S1 presents the average number of words generated in each specific category, for the three experimental tasks.

Next, we correlated the participants’ performance, quantified as the average number of words they produced in an experimental block, across the three tasks. Significant positive correlations in the participants’ performance were found between all three tasks, as shown in figure 3A (Pearson’s R and p-values displayed in the figure). Figure 3B depicts the distribution of the inter-event time intervals (IEIs) between consecutively produced words in the three experimental tasks. The IEI occurrence numbers were averaged across participants, using 2-second bins in the verbal fluency and instances task, and 4-second bins in the alternative uses task. The mean IEIs between consecutive word productions for each of the three tasks, averaged across the entire block duration, are also presented in figure 3B. The IEIs distribution showed a common trend in all 3 tasks, with a high frequency of short IEIs that decreases for longer IEIs. Furthermore, the IEI distributions fit a power law distribution, linearly fitting the log-log function, as shown in the insets in figure 3B. The vertical dashed lines in the plots mark the 12-seconds IEI cutoff for event-related analyses: only events that were preceded by an IEI that was at least 12 seconds or longer were further analyzed.

**Figure 3.**
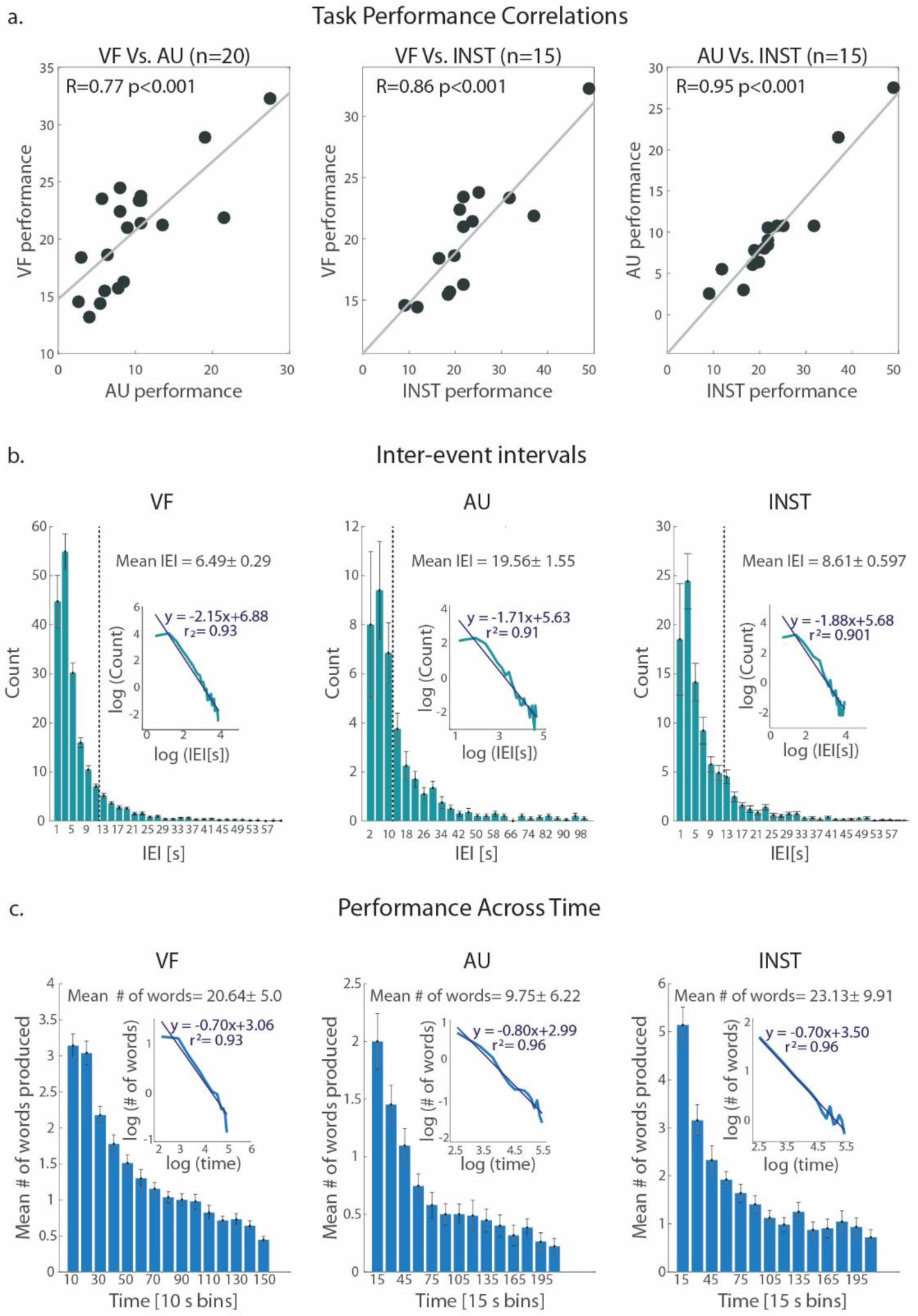
A. Correlations between the average number of words produced by individual participants in the three different tasks. Pearson R correlation coefficients and their p-values are specified. B. The IEI distribution, averaged across participants. Error bars denote the count average across participants± SE, for each IEI bin (2-second bins in the VF and INST plots, and 4-second bins in the AU plot). The total mean IEI value ±SE is indicated for each of the three tasks. The dashed lines indicate the 12-seconds mark, the IEI cutoff for further event-related analysis. The insets show the log-log plot and its’ linear fit for each task. C. Mean number of words produced during 10 or 15-second time bins (VF and AU/INST, in accordance) along the trial, averaged across all participants. Error bars denote the mean ±SE. The total mean number of words produced across the entire block duration ±SE is indicated for each of the three tasks. The insets show the log-log plot and its’ linear fit for each task.

In order to inspect the temporal dynamics of word generation across the block duration, we divided the blocks to 10-second (for the VF task) or 15-second (for the AU and INST tasks) time bins. Next, we counted the number of words generated in each bin, averaged across blocks for each participant, and finally averaged across participants. Interestingly, as demonstrated in figure 3C, there was remarkable similarity in the behavior across the three tasks, with an initially high word production rate, that decreased along block duration. As in the IEI distribution, the mean performance rate across time also followed the power law, linearly fitting a log-log function, shown in the insets in figure 3C. Together, these behavioral analyses reveal remarkable similarities across the three tasks, thus suggesting possible common underlying mechanisms.

### Anticipatory buildup in pupillary dilation

Previous studies have shown that pupil dilation is a reliable measure for task engagement and cognitive effort, even during covert behaviors (Einhauser et al., 2008; de Gee et al., 2014; Yellin et al., 2015; Broday-Dvir et al., 2018). Therefore, we measured eye movements and pupillary dilations during the three different experimental tasks, and compared the voluntary conditions to the externally driven control conditions. Average pupil size was locked to the word generation onsets, as indicated by the button press, for events that were separated by at least 5 seconds from the previous report, in order to avoid confounding signals from the previous event. Figure 4 depicts the pupillary dilation response during the free vs determined control events for the three tasks. A two-tailed paired t-test was calculated for each time point, comparing pupil size between the voluntary and the control conditions. As can be seen in the left panel, pupil size was significantly larger between times ∼ −1.7 seconds to ∼ 0 seconds relative to the word generation onset during the verbal fluency events, as compared to the control events, indicated by the yellow line (p<0.05, two-tailed paired t-test, fdr corrected). Similarly, pupil dilation was higher prior to the AU events, as compared to the control events, during the time period between ∼-2.8 seconds to −0.3 seconds preceding the verbal event generation (p<0.05, two-tailed paired t-test, fdr corrected). The INST task showed a similar pattern of higher activity ∼1 second prior to the voluntary events, as compared to the control events (p<0.05, two-tailed paired t-test, uncorrected), though the effect was weaker, likely due to the smaller number of participants with pupil measurements in this task. This task also showed a higher peak amplitude for the control response as compared to the instances mean response (p<0.05, two-tailed paired t-test, uncorrected), which could be related to the unpredicted sudden appearance of the external auditory cue (a similar trend, though insignificant, is seen in the AU vs. control responses).

**Figure 4.**
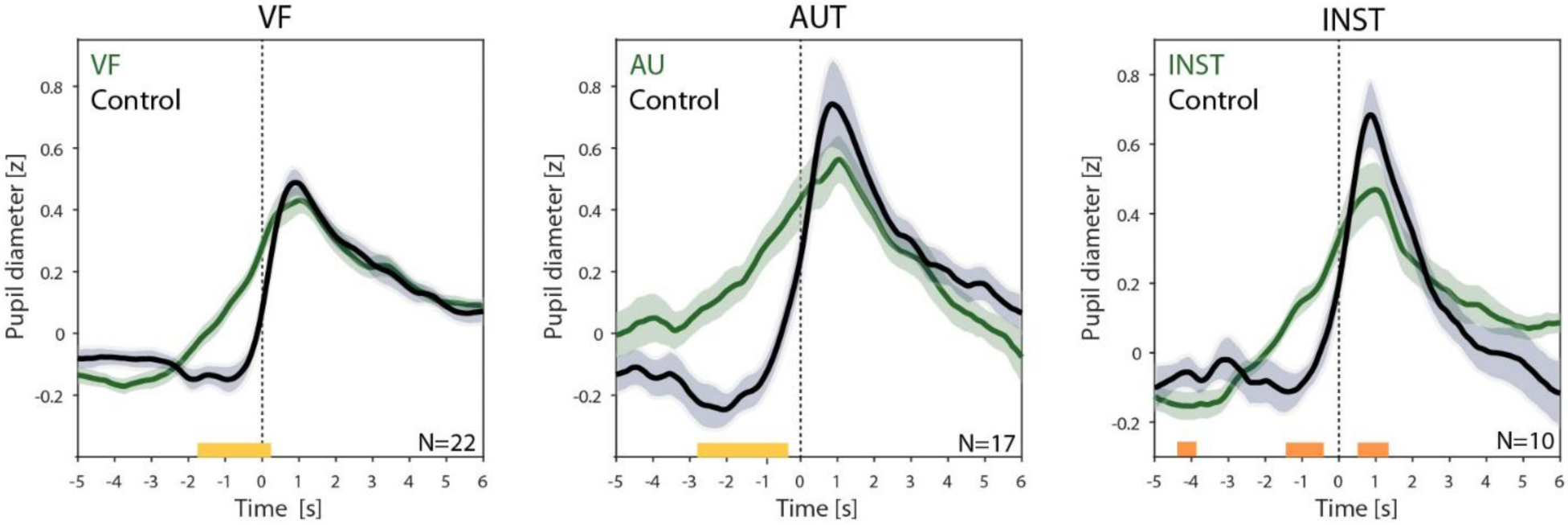
Group average pupil response during VF, AU and INST events and control verbal events, locked to the button presses. Transparent borders indicate the mean ± SE. Yellow lines indicate a significant difference between the two conditions (two-tailed paired t-test, p<0.05, fdr corrected). Orange line indicates p<0.05, uncorrected (though note the smaller number of participants in the INST task).

Thus, a 1-3 seconds slow anticipatory buildup was evident in pupil diameter preceding the voluntary, but not the externally-determined verbal events in all three experiments. This buildup likely reflects the increased processing and cognitive demand in the participants prior to the voluntary verbal generation, but not before the externally driven control events.

### Whole brain BOLD GLM results

Next we examined the BOLD-fMRI activations associated with the three tasks. Figure 5 depicts the whole brain GLM analysis contrasting the voluntary and controlled conditions for the verbal fluency (5a), alternative uses (5b) and instances tasks (5c). Unfolded cortices are presented on the left, showing the t-value group maps, with the pale-transparent colors reflecting the 0-threshold t-values, in order to allow full comparison between the three tasks. Overlaid, bright-colored and outlined activations, in warm colors (for increased activations) or cool colors (reflecting decreased activations) shown on the unfolded cortex, as well as on the inflated cortices on the right-hand side, depict the cluster corrected activations (voxel threshold of Z>2.3 and a corrected cluster significance threshold of p<0.05).

**Figure 5.**
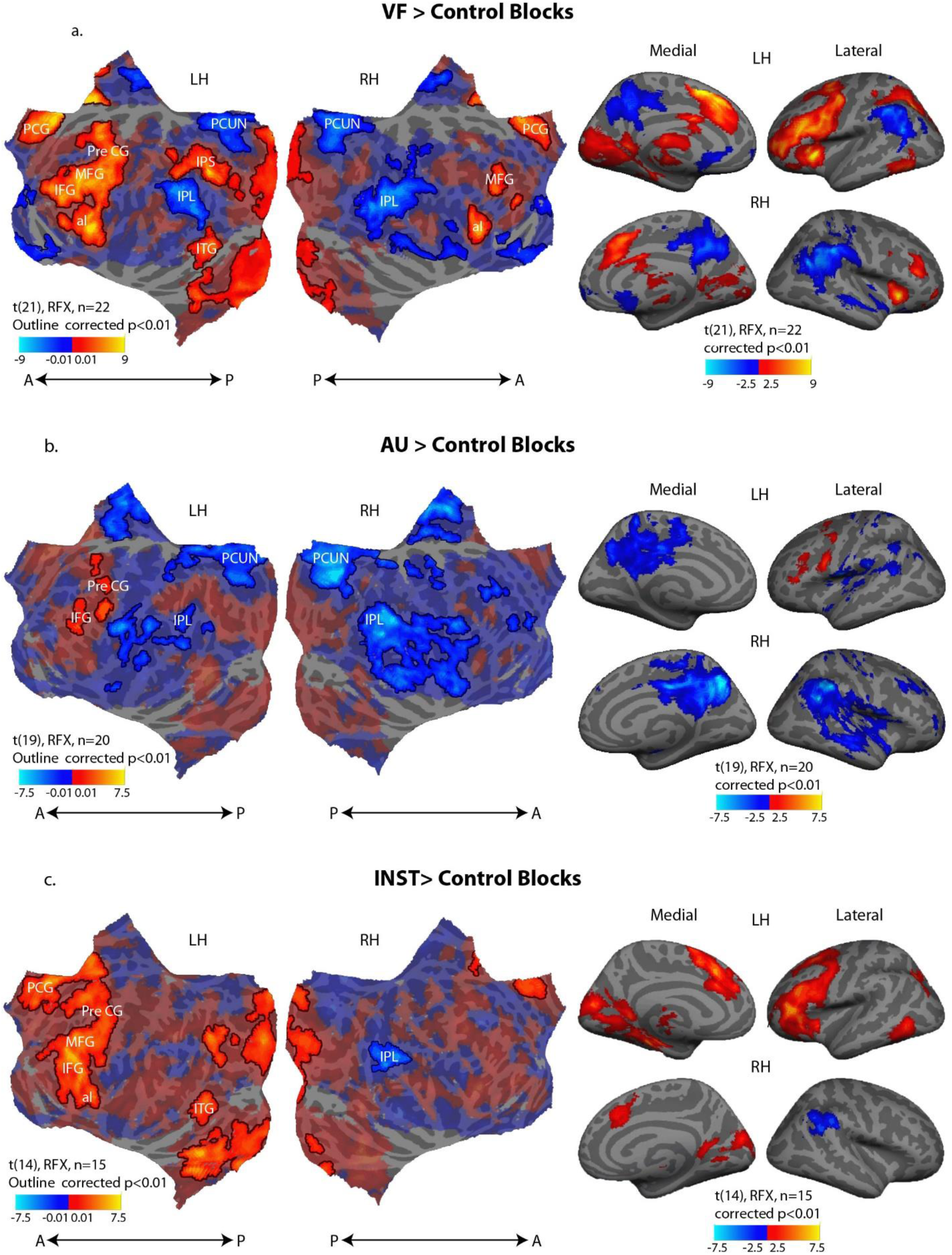
Whole brain t-value group contrast maps, generated by a random effects GLM analysis, for (A) Verbal fluency>control blocks, (B) Alternative uses >control blocks, and (C) Instances>control blocks. The activation maps are presented on top of inflated cortex (right panels, p<0.01, cluster corrected) and flattened cortex (left panels, pale-transparent colors show 0-thresholded data; outlined activations mark significant increases and decreases in activity, p<0.01 cluster corrected). Abbreviations: Paracingulate (PCG), Precentral gyrus (PreCG), middle frontal gyrus (MFG), inferior frontal gyrus (IFG), anterior insula (AI), inferior temporal gyrus (ITG), intra-parietal sulcus (IPS), precuneus (PCUN), inferior parietal lobule (IPL).

As can be seen, all three experimental tasks show increased BOLD activity during the voluntary blocks as compared to the control blocks in the left dorsal prefrontal cortex, specifically in the left inferior and middle frontal gyri (LH IFG and MFG), regions well-known to be involved in language production and processing. Increased activation was also apparent in the left precentral gyrus (PreCG), possibly related to the report of the generated verbal ideas by a button press. The anatomical distribution of these significant activations in the VF and INST tasks are relatively similar, while the extent of AU left prefrontal activation is more constrained. This could possibly be due to the significantly fewer ideas generated in AU blocks as compared to VF and INST blocks (see *behavioral results* section). Additionally, VF and INST blocks, as compared to control, showed significantly increased activations in the left anterior insula (AI), medial frontal regions including the paracingulate (PCG) and anterior cingulate cortex (ACC), the inferior temporal gyrus (ITG) and intra-parietal sulcus (IPS). AU blocks displayed a positive trend of activations in these regions as well, though they were much weaker and did not survive statistical corrections for multiple comparisons. These activations are in agreement with previous reports from VF and divergent and creative thinking imaging studies (e.g. Schlosser et al., 1998; Gonen-Yaacovi et al., 2013; Wagner et al., 2014; Wu et al., 2015).

In addition, the voluntary conditions, as compared to the control, showed significant decreases in default mode network (DMN) activity, specifically in the inferior parietal lobule (IPL) in all three tasks, and in the precuneus (PCUN) and posterior cingulate cortex (PCC) in the VF and AU tasks (with an insignificant trend in the INST task). Supplementary figure S2 depicts the averaged time-courses of the DMN responses to verbal events, for the IPL and PCUN ROIs. As can be seen, both DMN ROIs revealed consistent gradual decreases in BOLD activity 2-4 seconds prior to the onset of voluntary verbal events, but not before the externally-driven control events. As denoted by the yellow line, the DMN activity was significantly lower around the time of the voluntary event as compared to the control event, in all three tasks (two-tailed paired t-test, p<0.05, cluster correction). These DMN decreases might reflect the inhibition of mind-wondering or other irrelevant thoughts at the time of the production of a specific exemplar or creative idea.

### Anticipatory buildup in BOLD activity

In order to examine the dynamics of the fMRI-BOLD activations during the experimental tasks, we defined 4 frontal ROIs: the anterior part of the LH IFG (or pars triangularis), the posterior part of the LH IFG (or pars opercularis), the paracingulate cortex and the LH precentral gyrus. In order to avoid circularity, the ROIs were defined individually per participant based on the contrast of the first 30 seconds of the voluntary blocks (VF, AU or INST) > baseline (thresholded at p<0.0001, uncorrected), in conjunction with the anatomical masks of these regions, based on the Harvard-Oxford Cortical Structural Atlas (Desikan et al., 2006). An additional region, the LH anterior insula, was defined for the VF task only, but not for the other two tasks as it did not survive the statistical thresholding in most participants for these tasks. We then sampled the BOLD percent signal change time-course, locked to the time of the voluntary and control verbal generation events, excluding the events that occurred in the first 30 seconds. Furthermore, only events that were separated by 12 seconds from the previous event were considered, for both the experimental and the control conditions, in order to ensure that the inspected activity preceding the events was free from previous event-related activations (see *ROI definition and time-course extraction* and *ROI event-related response analysis* in *Online Methods*).

Figure 6 depicts the mean BOLD event-related activations in the anterior LH IFG (pars triangularis) ROI, outlined in cyan in the inflated cortices, for the three tasks, showing the voluntary responses (left panels), the control responses (middle panels), and the difference between them, defined as the within subject voluntary minus control time-course (right panels) differences, averaged across participants. Time points in which the response amplitude was significantly higher than baseline, defined as the mean amplitude across times −6 and −4 seconds before event onset, are denote by the yellow line (p<0.05, one-tailed paired t-test). Examining the VF response (figure 6A), it can be seen that the voluntary response showed a gradual build-up, reaching a significant increase above baseline at time 0, and preceding the control response, that reached significance only 4 seconds later. This trend was also seen in the differential averaged time-course, though reaching significance only later, at 2 seconds after event onset. Importantly, the fMRI time series were not shifted to compensate for the hemodynamic lag, so that the actual neural activity was increased ∼2-4 seconds earlier.

**Figure 6.**
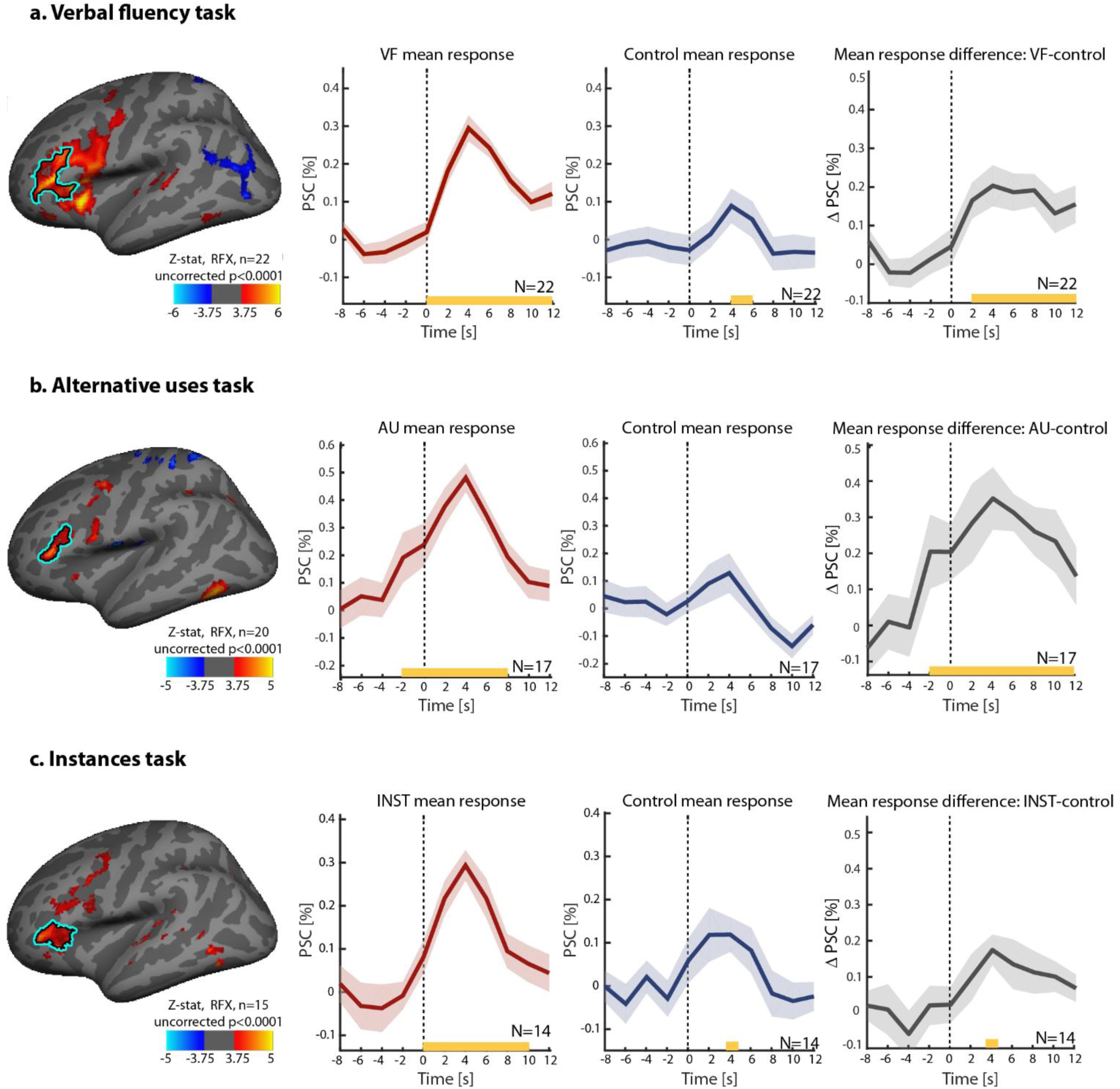
Group mean voluntary and control word generation responses, and their averaged response difference in the anterior LH IFG (pars triangularis), for: A. Verbal fluency (VF) task; B. Alternative uses (AU) task; and C. Instances (INST) task. Inflated left hemispheres display, for each task, the group contrast of the first 30 seconds of the voluntary blocks (VF/AU/INST)>baseline, thresholded at p<0.0001 uncorrected, shown for ROI definition illustration purposes. The anterior LH IFG ROI is denoted by the cyan outline. The ROIs were defined individually for each participant, based on this contrast at the single subject level in conjunction with the anatomical mask of the pars triangularis (see *ROI definition and time-course extraction* in *Online Methods* for details). Note the presented ROI outlines here are defined based on the group maps, solely for visualization purposes. Mean percent signal changes in the BOLD signal, averaged across participants, are presented in red for the voluntary conditions, blue for the control conditions, and gray for the mean response difference, defined as the voluntary event time-course minus the control time-course. Transparent borders indicate the mean ±SE. Dashed vertical lines indicate the time of the button press, reporting a verbal generation event (voluntary and control). Yellow lines indicate a significant increase in response amplitude above baseline, defined as the average amplitude across times −6 and −4 seconds before event onsets (one-tailed paired t-test, p<0.05).

Similar analyses were conducted for the AU (figure 6B) and INST (6C) tasks. As in the VF condition, the results revealed a significant anticipatory buildup of BOLD activation during the voluntary, but not during the control conditions. Specifically, the AU response reached a significant increase at time −2 seconds before the voluntary event onset, and the INST response at time 0. Increases in both voluntary responses preceded their corresponding control responses, evident also in the differential time-courses.

Similar dynamics were observed in the other ROIs studied. Supplementary figures 3,4 and 5 show the voluntary, control, and difference averaged time-courses for the VF experiment (figure S3), the AU experiment (figure S4) and the INST experiment (figure S5), for all the ROIs defined, specifically the posterior LH IFG (pars opercularis), paracingulate cortex, LH precentral gyrus, and the LH anterior insula for the VF task. The preceding anticipatory buildup was evident in all the task-related ROIs that were examined, prior to the voluntary verbal events, but not before the externally driven control events, in the three experimental tasks (except for the paracingulate in the AU task, in which there was no difference between the voluntary and control conditions in the time of the significant increase in amplitude). This suggests that the anticipatory buildup was a common effect in all ROIs participating in the voluntary task.

A possible confound that could lead to an apparent gradual buildup preceding the voluntary events, but not the control events, could be related to the peak amplitude difference between the two conditions. In order to control for this possibility, we normalized the responses of individual participants to their peak value, resulting in equal peak amplitudes of 1 in both the voluntary and the control responses. As shown in Supplementary figures S6, S7 and S8 (for the VF, AU and INST tasks, accordingly), the signal increases during the voluntary responses preceded the control related activations even after this normalization, demonstrating that the anticipatory buildup was not related to an amplitude difference between the conditions.

### The anticipatory buildup is not an artifact of time-jittered responses

Our results so far demonstrate a significant gradual buildup both in pupil dilation as well as in BOLD activity during the voluntary events and not during the control, externally determined, events. This gradual ramping of activity anticipated the voluntary events by 1-2 seconds. However, it is important to note that this gradual buildup was found after averaging across multiple individual events. Critically, it could be the case that the gradual buildup did not occur at the single trial level, but was rather artificially produced by inaccuracies in the timing of the participants’ reports of the occurrence of their freely generated verbal events. A simulated demonstration of how such an artificial buildup could be produced is shown in figure 7, which depicts the two possibilities. As can be seen in panel A, even if in individual trials the activity leading up to the creative or verbal generation event was abrupt, i.e. step-wise, a jitter in the report timing may have “smeared” the average of these step-wise events. Thus, when averaging across individual trials, a gradual buildup is apparent, as is demonstrated for the average neural activity and BOLD response simulation estimates in figure 7A. Importantly, the mean of such time-jittered step function trials is indistinguishable from the mean of true gradual buildup occurring on every single trial, as shown in figure 7B.

**Figure 7.**
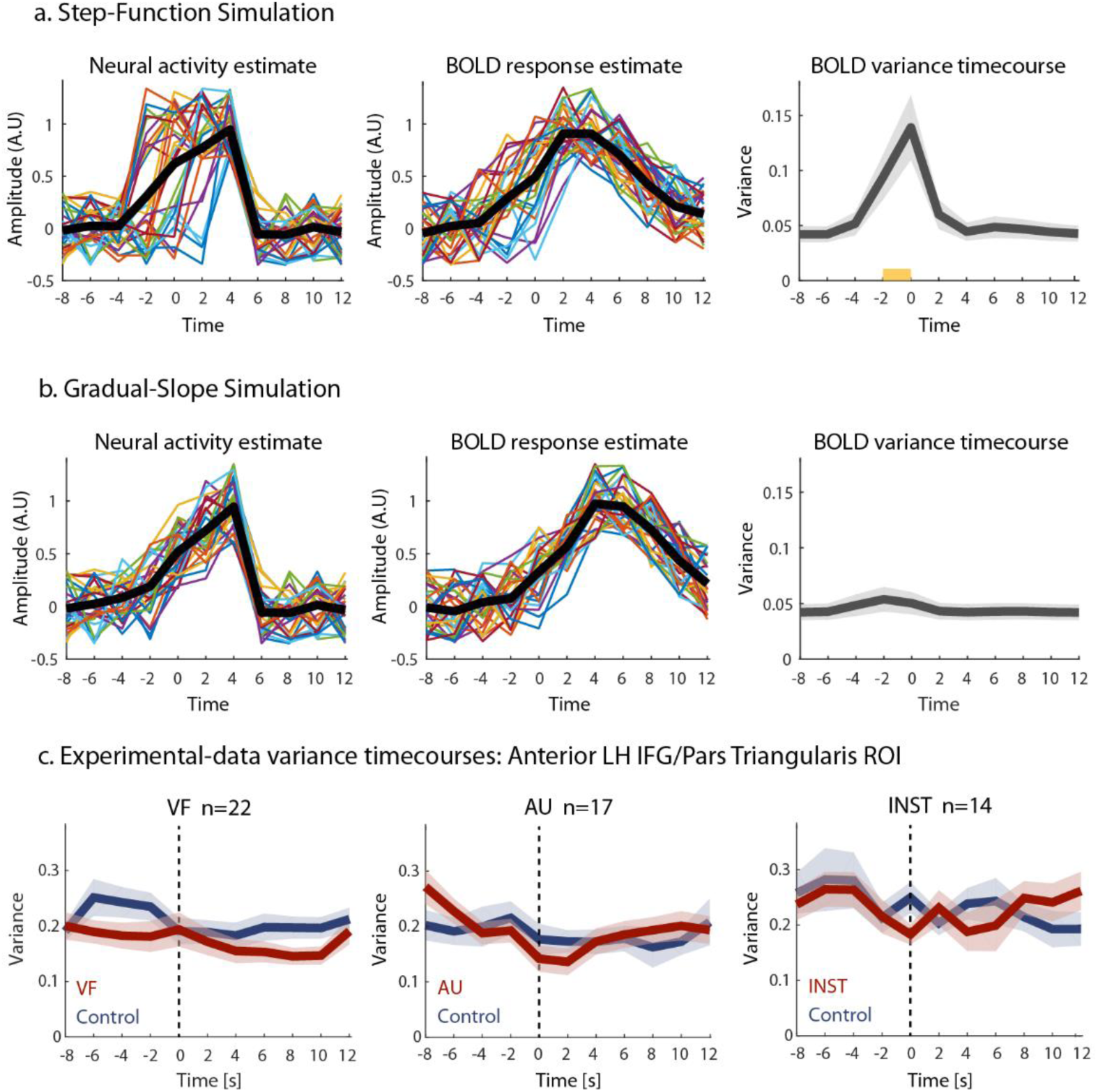
A. Step-function model simulation. The left panel displays the simulated neural activity of individual trials (n=30), shown in different colors. All trials begin at a baseline value of 0, rapidly increase to a peak amplitude of 1 during “1 TR” (2 seconds), and return to baseline 0 at time 6 s, with an addition of ±0.35 jittered noise to the signal. The time of the amplitude step-increase is jittered, occurring between times −2 to +4 s relative to “event onset”. The averaged time course across trials is shown in a thicker black line. The middle panel shows the BOLD response estimates of the individual trials, as well as the averaged signal, obtained through convolution of the neural activity simulation estimates with the standard HRF (Boynton et al., 1996). The left panel shows the time-course of the across-trials variance of the simulated BOLD responses, averaged across 1000 permutations. This averaged variance time course displays significant increases above baseline between −2 to 0 seconds relative to “event onset” (p<0.005, permutation test, baseline defined as the averaged variance at times −6 to −4 s). Transparent borders indicate the mean ±SE. B. Gradual-slope model simulation. Similar to (A), only here, instead of showing an abrupt step-increase from 0 to 1, all trials have a gradual positive slope of 0.25 ± 0.05 jitter, starting at −4 seconds with the value of 0 ±0.35 until reaching a peak of value 1 ±0.35 at 4 seconds. The simulated BOLD variance time course in this case is flat, with no significant changes. C. Variance time courses of experimental data from the anterior LH IFG ROI, in all 3 experiments, averaged across participants, for the voluntary conditions (red) and control conditions (blue). The time-courses are locked to event onset, as reported by button presses. In all tasks and conditions, the inter-trial variance remained relatively flat at all times, and did not significantly increase above the baseline variance, calculated as the average variance at times −6 and −4 seconds before event onset (one-tailed paired t-test, p>0.3 for all voluntary conditions at all time-points, and p>0.15 for all control conditions at all time-points). Transparent borders indicate the mean ±SE.

Fortunately, one parameter that can clearly differentiate between jittered step-wise function trials and true single-trial ramping is the across-trial variance. Thus, as was revealed in a simulation analysis, the time course of the inter-trial variance in the case of individual jittered step-functions shows a significant peak coinciding with the time of the anticipatory buildup, as shown in the right panel of figure 7A (p<0.005 at time points −2 and 0 seconds, calculated across 1000 permutations; values compared to the average variance at times −6 and −4). By contrast, single trial gradual-ramping stimulation displayed a flat across-trial variance time course (p>0.25 for all time points, across 1000 permutations), see right panel in figure 7B (for additional details regarding the simulation procedure, see *Variance control simulation and analysis* in *Online Methods*).

Panel C in figure 7 displays the time-courses of the across-trial variance of the experimental data from the anterior LH IFG, for the voluntary and control conditions separately. All three tasks revealed a relatively flat variance time-courses, with no significant increases during the anticipatory buildup duration, relative to baseline variance (p>0.3 for all time points in the voluntary conditions; p>0.15 for all time points in the control conditions; one-tailed paired t-test). The figure displays the variance from one example ROI in all three tasks, yet examining the additional ROIs defined, as well as the pupillary across-trials variance time-courses, revealed the same result, with no significant increases in variance during the duration of the gradual signal increase. Thus, our variance analysis clearly ruled out the possibility that the observed anticipatory buildup was due to time jitter of step-function-like, or extremely rapid, activations.

### Resting state fluctuations are correlated to the anticipatory buildup

Finally, we examined the critical question: was there a link between the resting state BOLD fluctuations, measured in a resting-state scan prior to the experimental task runs, and the anticipatory buildup preceding the free verbal and creative events? In order to examine this question, we employed the commonly used measure of the fractional amplitude of low frequency fluctuations (fALFF), a method that allows measuring the relative portion of slow frequency fluctuations from the entire detectable power spectrum (Zou et al., 2008; Zuo et al., 2010) (see *fALFF analysis and simulation* in *Online Methods)*. Furthermore, as shown in supplementary figure S9, fALFF values calculated from simulated “event-related” responses, as well as simulated “resting state time courses” derived from concatenated single “events”, are correlated with the slopes of these responses and time-courses, suggesting that this measure holds information regarding the shape of these responses or fluctuations. Furthermore, the fALFF values of simulated “event-related responses” and “resting state time courses” that share a common slope, also show a significant positive correlation, as shown in figure S9, further emphasizing the logic behind the usage of this common method for uncovering a link between the experimental resting state fluctuations and event-related responses (for additional details see *fALFF analysis and simulation* in *Online Methods*).

Therefore, we calculated the fALFF values of each individual participants’ resting state activity fluctuations, as well as the fALFF values of the mean VF, AU and INST voluntary event-related responses. The fALFF values of the matched average deterministic control responses from the three tasks were also obtained. To examine the possible link between resting state fluctuations and the anticipatory buildups, we correlated the fALFF values of the resting state fluctuations with the fALFF values of the voluntary condition responses, and compared them to the correlations between the resting state and the control response activations, across participants. The results of this analysis for all three tasks, from the anterior LH IFG (pars triangularis) ROI, are shown in figure 8. Left-side scatter plots depict the correlation between individual participants’ voluntary event-related responses and the resting state time courses, and the right-side plots show the correlations between the control activations and the resting state fluctuations (VF task results shown in A; AU task shown in B; and INST task in C). As can be seen, significant correlations were found between the voluntary average buildups and the resting state fALFF values, in all three tasks (VF task: Spearman’s R=0.71, p<0.001; AU task: Spearman’s R=0.52 p=0.017; INST task: Spearman’s R=0.62 p=0.009. All p-values derived from a subject-wise label-shuffling permutation test, across 10,000 permutations, and survived FDR correction for multiple comparisons at p<0.05). Critically, no significant correlations were found between the fALFF values of the mean deterministic control responses and the resting state time courses (VF task: Spearman’s R=0.10, p=0.32; AU task: Spearman’s R=-0.11 p=0.65; INST task: Spearman’s R=-0.05 p=0.56. All p-values derived from a subject-wise label-shuffling permutation test, across 10,000 permutations). To directly compare these correlations between the anticipatory buildup during voluntary events and the resting state dynamics to the correlations between the control activations and resting state, separately for each of the three tasks, we used a percentile bootstrapping test for comparing robust dependent correlations (Wilcox, 2016). This analysis revealed that all voluntary vs. resting state correlations were indeed significantly higher than their counterpart control vs. resting state correlations (VF task: p=0.02; AU task: p=0.032; INST task: p=0.008; dependent correlations test, one-sided α=0.05).

**Figure 8.**
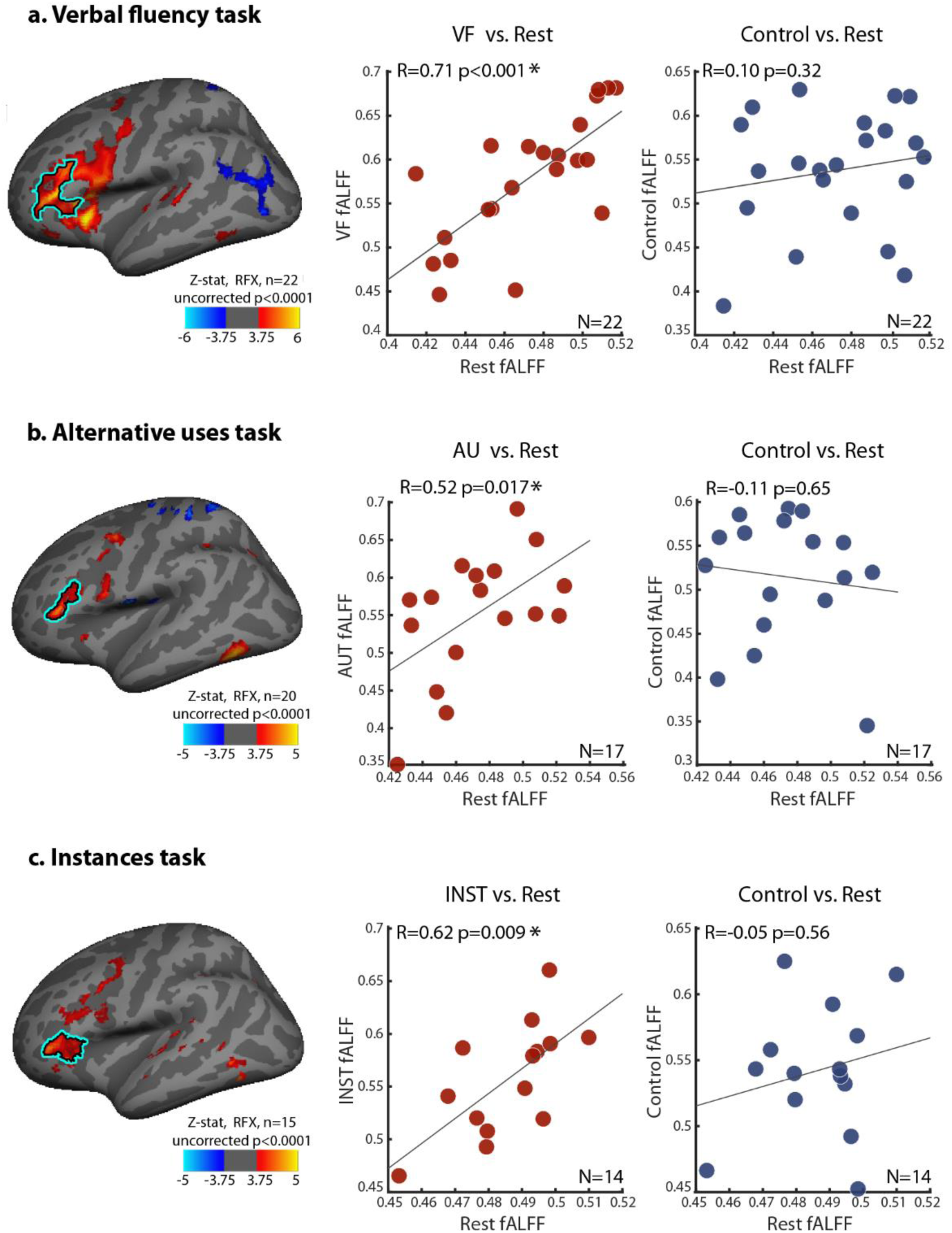
fALFF correlation plots, depicting the correlations between the fALFF values of participants’ mean voluntary responses vs. the fALFF values of individual participants’ resting state time-courses in the anterior LH IFG ROI/pars triangularis (shown in red markers, each marker specifies an individual participant); and the correlations between fALFF values of participants’ mean control activations vs. the fALFFs of their resting state time-courses (shown in blue markers), for the same ROI. The results are shown for the VF task (A), the AU task (B) and the INST task (C). The inflated left hemispheres presented here denote the anterior LH IFG ROI, and are identical to those in figure 6 (see figure 6 legend for details). Spearman’s R correlation coefficients are presented in the plots, together with their p-values, derived from a subject-wise label-shuffling permutation test (10,000 permutations). Significant correlations are marked with an asterisk (p<0.05, FDR corrected for multiple comparisons across the additional ROIs defined, presented in Supplementary figures S9, S10 and S11). All tasks showed significantly higher correlations between the voluntary buildups and resting state fALFFs, than between the control activations and the resting state fALFFs (p<0.05, dependent correlation percentile bootstrapping test (Wilcox, 2016)).

The voluntary vs. rest and control vs. rest fALFF across-subject correlations were inspected in additional ROIs, aside from the anterior LH IFG, including the posterior LH IFG (or pars opercularis), paracingulate cortex, LH precentral gyrus and LH anterior insula (for the VF task only). The correlation plots for these ROIs are presented in the supplementary figures S10 (for the VF task), S11 (AU task) and S12 (INST task). Significant correlations between the fALFF values of mean voluntary responses and resting-state fluctuations were found in the posterior LH IFG ROI in the VF and AU tasks (VF task: Spearman’s R=0.58, p=0.002; AU task: Spearman’s R=0.57 p=0.006; All p-values derived from a subject-wise label-shuffling permutation test, across 10,000 permutations, and survive FDR correction at p<0.05), and these correlations were higher than the non-significant correlations between control and resting-state in this ROI (VF task: p=0.012; AU task: p=0.04; Wilcox dependent correlations test, one-sided α=0.05, statistical significance marked by yellow frames in figure S10 and S11). Yet this effect in the posterior LH IFG was not evident in the instances task (Spearman’s R=0.02, p=0.47, permutation test). Additional positive correlations were found between the AU voluntary response and resting state in the paracingulate cortex (Spearman’s R=0.68, p<0.001, permutation test, survived FDR correction at p<0.05) and between the INST voluntary response and rest in the LH precentral gyrus (Spearman’s R=0.64, p=0. 008, permutation test, survived FDR correction at p<0.05), though in both cases, the correlations were not statistically higher than the correlations between control and rest (p=0.12 and p=0.31, accordingly, Wilcox dependent correlations test, one-sided α=0.05). Importantly, no significant correlations were found between the mean control responses and resting state fluctuations, in all tasks and in all ROIs examined (see figures S10, S11 and S12 for all correlation plots and R and p values).

Thus, our analysis revealed a significant correlation between the dynamics of the anticipatory build-up preceding voluntary verbal events, and the dynamics of resting state fluctuations. Critically, this correlation did not exist between the externally-driven control verbal events and resting state, thus ruling out that this effect is possibly a general BOLD-related or language-production related phenomenon. Rather, it indicates that the link between resting state and task was specific to the free, inner-generation mode. The significant voluntary-rest correlations were apparent in all three voluntary tasks, suggesting a common mechanism for free language production, creative verbal production and verbal divergent thinking. This effect was also specific to the left IFG ROIs, regions that are well known to be involved in language processing and production (e.g. Petersen et al., 1988; Gabrieli et al., 1998).

## Discussion

Our study supports the notion that a common neuronal mechanism underlies all types of free voluntary behaviors. In our study we focused specifically on voluntary verbal behaviors, including the free generation of verbal exemplars, ideas and creative thoughts. By using three different tasks-a very common language production and fluency task (verbal fluency or VF), a classic verbal creativity task (alternative uses or AU) and a verbal divergent thinking task (instances or INST), we were able to highlight the common neuronal mechanism for the internal, unpredictable, “free” generation of a verbal idea, that can be generalized across different tasks and contexts.

Our findings thus extend previous studies (reviewed in Moutard et al., 2015) indicating that a common neuronal “signature” of free behavior (as we operationally defined in the introduction) is a slow buildup of activity in the relevant task-related networks preceding the actual moment of free behavior. In the present paper we have extended this common principle to the case of free verbal behaviors. Specifically, in all three tasks, we found a slow, gradual buildup of BOLD signal preceding the reported time of the freely-generated verbal idea or creative event by ∼1-2 seconds, evident in language and additional task-relevant brain regions (see figure 6). Importantly, this gradual anticipatory increase was not present before the control events: deterministic, externally-driven word repetitions. Thus, the anticipatory buildup appears to be specifically linked to free, voluntary, internally-generated events, and not to verbal responses in general.

Further support for the validity of the anticipatory buildup as a signature of free behavior is provided by our pupil dilation measures. These pupillary measurements were conducted concurrently with the task performance and manifested a slow buildup of similar dynamics prior to the voluntary “free” events, but not before the deterministic control events (as shown in figure 4). This pupil dilation is in concordance with previous works, by ourselves and others, that have proposed that such pupil dilations provide a reliable index to processing levels (e.g. Kahneman & Beatty, 1966; Alnaes et al., 2014; de Gee et al., 2014; Yellin et al., 2015; Broday-Dvir et al., 2018), and this observation was nicely compatible with the suggestion that free behaviors are characterized by a slow anticipatory buildup of subliminal activity.

An important concern to be ruled out is that the observed gradual ramping of activity was in fact merely an artifact, caused by the lack of precision in participants’ ability to accurately report the timing of the emergence of the verbal idea or exemplar. Thus, it could be argued that in fact each free-behavior event is characterized by a rapid, step-wise, activation increase (see figure 7). Under such an interpretation, the gradual buildup that was observed might in fact be merely a “smeared” byproduct of averaging multiple step-functions with jittered timings. Indeed, in our simulation analysis (presented in figure 7), we were able to recreate a slow buildup by averaging a set of such temporally-jittered step-function responses. However, a major discrepancy between the true, single trial, gradual buildup model we proposed, and the averaged, jittered step-function model, is revealed when considering the inter-trial variance (right-side panels in figure 7A and B). Here, the jittered step-function model predicts a significant increase in the cross-trial signal variance during the buildup period (due to the jittered event timing), while the single trial-ramping model predicts similar variance levels across the entire anticipatory period. Careful examination of the BOLD variance during our experiments revealed a flat variance time-course, with no increases (or any significant differences at all) compared to baseline variance levels, unequivocally supporting a real gradual buildup prior to every voluntary event (see figure 7C).

What could be the neural mechanism that accounts for the slow anticipatory buildup? Two aspects of this phenomenon are helpful in narrowing the range of possibilities. First, free behavior is an extremely ubiquitous phenomenon, occurring at diverse modalities and functions (Moutard et al., 2015), from the classically studied decisions to perform a simple movement (Libet et al., 1983; Schurger et al., 2012), to spontaneous music and dance improvisation, as well as idea generation and creative behaviors. Even tasks that are typically considered to be mainly stimulus driven, such as visual perception, can manifest free or voluntary aspects, for example in the spontaneous alterations during bistable perception (Gelbard-Sagiv et al., 2018) or spontaneous visual imagery (Norman et al., 2017; Norman et al., 2019). Consequently, if our hypothesis states that there is a shared neural mechanism underlying this diverse set of free behaviors, then it should be a ubiquitous neuronal process, that can be found essentially across all cortical networks.

Another key aspect of this neuronal process is its slow dynamics: examining both the BOLD activity changes and the pupillary dilations here, as well as EEG and ECoG signals from previous studies of free behavior (Gelbard-Sagiv et al., 2008; Schurger et al., 2012; Norman et al., 2019), reveals a process that has a time constant of 1-2 seconds. This is an order of magnitude slower than the typical 200-millisecond stimulus-response cortical dynamics (e.g. Bitterman et al., 2008; Fisch et al., 2009; Podvalny et al., 2017).

Examining the possible neuronal candidates manifesting both slow and ubiquitous cortical activity reveals a readily obvious candidate: the spontaneous (also termed resting-state) fluctuations that have been observed and studied extensively across essentially every human cortical network (Biswal et al., 1995; Arieli et al., 1996; Nir et al., 2006; Fox & Raichle, 2007). So how can these ultra-slow resting state fluctuations account for the ramping buildup observed prior to free behaviors? Our hypothesis, extended from an earlier proposal by Schurger (Schurger et al., 2012) for the case of volitional movements, is illustrated in figure 1. Essentially, we propose that free and creative behaviors are initiated by the slow resting state fluctuations. These fluctuations drive the network that is relevant for the voluntary task across a decision threshold. Thus, prior to free behavior, the activity in the network manifests slow spontaneous fluctuations, and when such a fluctuation crosses the activation threshold, a free behavior can emerge. A strong prediction of this hypothesis is that prior to each and every free behavior event, we expect to see the slow uprising phase of a spontaneous fluctuation, hence the slow ramping activity evident prior to the free verbal and creative moments shown here.

An obvious counter-argument to this proposed mechanism could be that the observed similarity between the slow time constants of the resting state and the anticipatory slow buildup are a mere coincidence. Accordingly, it could be argued that the two phenomena are unrelated, and are derived from completely different mechanisms that simply happen to both exhibit slow time constants.

If this was indeed the case, the temporal dynamics of the resting state fluctuations and the anticipatory buildup should also be independent of each other. However, measuring the response characteristics across individuals revealed significant positive correlations between the fractional amplitude of low frequency fluctuations (fALFF) of the resting state time-courses and the anticipatory buildup preceding the freely generated verbal responses (see figure 8). This correlation was evident across all three different tasks that were examined, including the verbal fluency task-a classic language production task (Troyer et al., 1997; Schlosser et al., 1998), the alternative uses task-used commonly for verbal creativity assessment (Guilford, 1967; Torrance, 1988), and the instances task, a divergent thinking task (Silvia, 2011). This replication across the three separate tasks suggests that the resting state fluctuations constitute a common mechanism underlying a variety of diverse voluntary verbal behaviors. The correlation between the anticipatory buildup and the resting state fluctuations was spatially specific to the LH IFG, the central region involved in language processing and generation, which are of course the key components of these tasks. Importantly, the correlations were specific to the free-behaviors, and were not evident for the deterministic externally-driven control tasks (see figure 8). Thus, the correlations could not be attributed to general individual differences in the BOLD responses, or to general verbal-related or language-related responses, but rather were specific to voluntary, internally-driven events.

Together, these results strongly support our hypothesis (shown in figure 1A) that the resting state fluctuations constitute a common neuronal mechanism that drives free behaviors, and that their rising phase constitutes the anticipatory buildup observed prior to the initiation of these behaviors.

Our study strongly supports the notion that creative behaviors rely on a similar neuronal mechanism that drives free verbal behaviors in general. Three aspects of our results support this conclusion. First, our behavioral results show significant similarities in the participants’ performance across the three different tasks, including correlations in number of words and verbal ideas produced, as well as common temporal dynamics of production, shown in figure 3. Second, the slow anticipatory buildup, both in the BOLD signal as well as in the pupillary response, precedes all voluntary events, including the fluency and creativity tasks, but not the control events. Finally, the link between resting state fluctuations and the anticipatory buildup was evident in the three different tasks as well. Additionally, the proposed role of spontaneous fluctuations in the emergence of creative thoughts and verbal ideas also fits nicely with previous reports of links between resting state activity patterns and creative abilities (Takeuchi et al., 2012; Beaty et al., 2014; Yin et al., 2015; Beaty et al., 2018; Shi et al., 2019; Sun et al., 2019), as well as changes in resting-state connectivity patterns following creative or divergent-thinking training (Wei et al., 2014; Fink et al., 2018).

Of course, the verbal exemplars or creative ideas that are generated depend on previous experience and learning, the underlying brain structure and connectivity, etc. Here we suggest that the generation process, in which a new idea is created, or a specific exemplar comes to mind (out of a dozen or more relevant possibilities), involves accumulation of stochastic noise in the specific task-relevant region, in our case the language-related ROI. We propose that the spontaneous, or resting-state fluctuations, constitute a stochastic exploration process, that may tilt the system towards a specific idea, exemplar or creative thought at a random moment in time.

Given the importance of creative behavior to human progress and achievements, our findings, pointing to resting state fluctuations as a candidate driving mechanism for creativity, are particularly significant. It is perhaps no coincidence that many anecdotal reports of creative ideas and problem solving in day-to-day life, as well as stories of great ideas or inventions, occur while taking a walk (Nikola Tesla and the AC motor), in the shower or bath (Archimedes’ principles of density and buoyancy) or while daydreaming in public transport (J.K Rowling and Harry Potter). These are all situations that reduce attention to specific tasks or external stimuli, and hence likely enhance resting-state fluctuations. As our results suggest that these intrinsic fluctuations are involved in driving idea generation, it is an intriguing question whether this type of behavior will be enhanced in contexts in which these fluctuations flourish. Thus, our present findings may open future informed directions for identifying the optimal conditions and even developing methods for enhancing human creativity.

## Online Methods

### Participants

In total, 24 healthy, right-handed participants (11 female, mean age 27.56± 3.89) with normal vision participated in the study, that included three different tasks in two separate scanning sessions. One participant was excluded due to excessive motion. Another participant was excluded from one scanning session due to excessive motion, while his second session was maintained. All participants provided written consent and received payment for their participation. All procedures were approved by the local ethics committee. The experiment included three different tasks: a verbal fluency (VF) task, an alternative uses (AU) task and an instances of common concepts (INST) task. 22 participants participated in the VF task (10 female, mean age 27.5±3.97), 20 in the AU task (10 female, mean age 27.7± 4.16) and 15 in the INST task (8 female, mean age 27.47± 4.75). 15 participants completed all three tasks, 4 completed only the VF and AU tasks, 3 completed only the VF and 1 completed only the AU.

### The experimental tasks and design

Our study included a total of three different verbal tasks: a verbal fluency (VF) task, an alternative uses (AU) task and an instances (INST) task, that were completed in 2 separate scanning sessions. One session included 5 runs of the VF task, and the other session included 2 AU runs and 2 INST runs. Session and run order were counterbalanced and randomized. All experimental sessions were initiated with a short training period outside magnet, in order to familiarize the participants with the task/tasks that were to be performed in the current scanning session. Next, the participants entered the scanner and underwent an 8-minute resting state scan with eyes open. Then, the participants either completed 5 VF task runs, or 2 AU and 2 INST runs. All tasks were performed in a covert manner, in order to avoid movement-related artifacts due to overt speech, specifically artifacts during the time period preceding to the idea onset and report. An anatomical scan was also completed after 2 experimental runs. Figure 2 depicts the three tasks as well as the control condition, which was the same in all experimental runs.

Each VF block was initiated with a visual cue instructing the participants to covertly generate words from a specific category, that could be either a phonemic or a semantic category. The semantic categories were tools, birds, fish, vegetables and USA states; the phonemic categories were words that begin with the Hebrew letters “yud”, “tzadik”, “pay”, “vav” and “tet”. The semantic categories were intended to be relatively harder than more general categories, such as animals or food and drinks, in order to try to induce a slower generation pattern, that will lead to longer time intervals between the words generated. The participants were instructed to press a button immediately every time they thought of a new word from the relevant category, after which a short audio “beep” was heard. Each VF block lasted 2.5 minutes and was terminated by the visual cue “Break”, followed by a 30-second break period.

Control blocks began with the visual instruction “repeat the word ‘__’ when the auditory cue is heard”, with a different specified word to be repeated on each block. Here, the participants needed to wait until they heard the auditory cue. Once it was played, they were instructed to covertly repeat the instructed word, and immediately afterwards press the button. Importantly, the auditory cues in the control blocks replayed the participants’ performance in the VF blocks, specifically reconstructing the participants’ inter-response intervals during the 100 final seconds of these blocks. This manipulation allowed us to compare similarly spaced control and VF events in during the data analysis stages. Control blocks were 100 seconds long, and were also terminated with the visual cue “Break”, followed by a 30-second break. Each experimental run lasted ∼10 minutes, and included 2 VF blocks, one of a semantic category and one phonemic, and two control blocks, in random order. In both the control and VF conditions, the initial visual instructions for each block appeared for 2 seconds, after which a blank gray screen was presented for the rest of the block. Participants were instructed to maintain their gaze in the middle of the screen for the entire block duration.

The AU task, a divergent thinking task used commonly in creative thinking studies, requires the participants to think of creative uses for everyday objects: a button, a paperclip, a barrel and a cup (Guilford, 1950; Arden et al., 2010; Dietrich & Kanso, 2010; Beaty et al., 2014). AU blocks began with the instruction to generate creative uses for a specific object. Participants then needed to press the button every time a new idea came to their mind, after which the auditory “beep” was played. AU blocks were terminated after 210 seconds with the cue “Break”, followed by a 30-second break period. Control blocks were similar to those in the VF experiment described above, requiring “passive” word repetition that replayed the participants’ AU block performance, with 120 seconds long blocks. Each AU run lasted ∼ 13 minutes, and included 2 AU blocks and 2 control blocks, in random order. The final task, the INST task, had a similar structure to the two previous tasks, only here the participants needed to generate exemplars for common instances, including things that are heavy, loud, round and tall (Silvia, 2011). INST blocks were 210 seconds long, during which the participants pressed a button each time they thought of a new exemplar. Control blocks, same as in the other tasks, lasted 120 seconds. INST runs were ∼ 13 minutes, and included 2 INST blocks and 2 control blocks.

By setting long voluntary-block durations, in all three tasks, we were able to obtain voluntary and control verbal events that were separated in time with a relatively long “no-event” period preceding them. Specifically, only events that were separated by at least 12 seconds from the previous event were further examined in the event-related analyses. This allowed us to inspect the “clean” neural activity leading up to the subjective experience of a spontaneous emergence of an idea, as compared to the control passive word production in response to an external cue, without contamination of the previous event-related response.

At the end of each experimental session, the participants filled out questionnaires reporting the all the words and creative ideas they remembered generating for each category that was probed. This was done in order to ensure they had understood the tasks correctly, and generally sample their responses.

### MRI setup

16 participants (of which all 16 completed the VF experiment, and 15 completed the AU and the INST experiments) were scanned in the 3 Tesla MRI scanner (Magnetom Prisma, Siemens), at the Weizmann Institute of Science, using a 20-channel receive head/neck coil. Functional images of blood oxygenation level dependent (BOLD) contrast were obtained using a T2* -weighted gradient echo planar imaging (EPI) sequence (TR =2000 ms, TE = 30 ms, flip angle = 75°, FOV 210 mm, voxel size 3.0×3.0×3.3 mm, 32 slices, tilted to the ACPC plane). Whole-brain T1-weighted anatomical images were acquired for each participant using a 3D Magnetization Prepared Rapid Acquisition Gradient Echo (MPRAGE) sequence (TR = 2300 ms, TE = 2.32 ms, TI = 900 ms, flip angle = 8°, voxel size 0.9×0.9 ×0.9 mm, with integrated parallel acquisition (iPAT) acceleration factor of 2).

The additional seven participants (of which 6 completed the VF experiment and 5 the AU experiment), were scanned in the 3 Tesla MRI scanner (Tim Trio, Siemens) at the Weizmann Institute of Science, using a 12-channels head matrix coil. Functional images of blood oxygenation level dependent (BOLD) contrast were obtained using a T2*-weighted gradient echo planar imaging (EPI) sequence (TR =2000 ms, TE = 30 ms, flip angle = 75°, FOV 216 mm, voxel size 3×3×3 mm, 32 slices, tilted to the ACPC plane). Whole-brain T1-weighted anatomical images were acquired for each participant using a 3D MPRAGE sequence (TR = 2300 ms, TE = 2.98 ms, TI = 900 ms, flip angle = 9°, voxel size 1×1 ×1 mm).

### Pupil size and eye-tracking acquisition, preprocessing and analysis

Pupil size and eye-movements of the participants’ dominant eye were recorded continuously throughout the experimental scanning sessions, using an Eyelink-1000 eye-tracking device (SR Research, Osgoode, ON, Canada), at a sampling rate of 500 Hz. The pupillary data of 3 participants was not recorded in the AU and INST sessions, due to technical issues. Furthermore, scans in which the amount of missing data, due to technical issues or excessive blinking, was larger than 20% were also removed, resulting in the exclusion of two additional participants from the INST task analysis. In total, pupillary analyses included 22 participants for the VF task, 17 participants in the AU task, and 10 participants in the INST task.

The scaled index estimate of the pupil diameter, as recorded by the Eyelink system, was preprocessed using custom-made MATLAB scripts, following previously reported pipelines (Yellin et al., 2015; Broday-Dvir et al., 2018). Points of missing pupil data or unlikely pupil size (3 SDs from mean pupil size within a trial) were removed, along with their neighboring data points 80 ms before the onset and after the offset of the detected segments. The resulting gaps of missing data were replaced using an inverse-distance weighted interpolation (Howat, University of Washington, 2007). The entire pupillary time-course of each run was then bandpass filtered, removing frequencies outside the 0.015-5 Hz range, using the least-squares FIR filter adapted from the EEGLab MATLAB toolbox (Delorme & Makeig, 2004). Next, the time-courses underwent z-score normalization.

Pupil event-related response onsets were defined as the moment of the button press, by which the participants reported generating a new verbal exemplar or idea in the voluntary conditions, or repeating a specific word in the control condition. The pupillary time-courses for each event were extracted from −5 seconds before event onset, to 6 seconds after the event onset. Only creative and control events that were preceded by at least a 5-seconds event-free period were further analyzed, while the events that were not “isolated” in time, as defined, were discarded.

In order to rule out eye-movements as possible confounds to pupil dilation differences between the voluntary and the control conditions, we ensured no eye-movement differences were found between the two conditions. This was done by first calculating the Euclidean distance of the eye gaze location from the screen center, as determined by the x and y gaze coordinates, for each time point in each trial. Next, the mean distance and the variance of the distances from screen center were calculated per trial, and averaged across voluntary and control trials separately for each participant, for each of the three experiments. For the VF experiment and the INST experiment, no significant differences were found in the mean eye-movement distance between the voluntary and control conditions (paired two-tailed t-test, VF experiment, for VF>control: t(21)= −0.645, p= 0.526; INST experiment, for INST>control: t(9) = 0.158, p=0.88). Similarly, no significant differences were found in the variance of eye-movements between the voluntary and control conditions in these two tasks (paired two-tailed t-test, VF experiment, for VF>control: t(21)= 0.532, p= 0.60; INST experiment, for INST>control: t(9) = 1.83, p=0.10). The AU task, however, did show significant difference in the eye-movement distance mean and variance between the two conditions (p<0.05). In order to control for this, we removed both voluntary and control trials in which the mean distance or mean distance-variance of that trial were larger by 2 s.d. or more from the mean values across trials of each participant. These trials were discarded from further analyses. Following this procedure, the mean and variance of the eye-movement distances were not different between the voluntary and control conditions in the AU task as well (paired two-tailed t-test; for mean AU distance>mean control distance: t(16)= 1.583, p= 0.133; for AU distance variance>control distance variance: t(16)= 1.581, p= 0.134).

Next, pupil event-related responses were averaged across individual trials in each participant, separately for the voluntary and the control conditions in each of the three experiments, and the responses were then averaged across all participants. In order to examine the difference between the voluntary and control conditions, separately in each experiment, we used a paired two-tailed t-test at each time point, and corrected for multiple comparisons using FDR correction, according to Benjamini-Hochberg method (α= 0.05) (Benjamini & Yekutieli, 2001).

### fMRI preprocessing

MRI data processing was achieved using FSL 5.0.4 (https://fsl.fmrib.ox.ac.uk/fsl/fslwiki/) and in house MATLAB codes (R2016b, The MathWorks). The functional task data was preprocessed using FEAT version 6 (FMRIB’s expert analysis tool), and included the following steps: removal of the first 2 volumes from each functional scan; motion correction using FMRIB’s Linear Image Registration Tool MCFLIRT (Jenkinson et al., 2002); brain extraction using BET (Smith, 2002); high pass temporal filtering of 100s; and spatial smoothing using a Gaussian kernel of FWHM=6mm. Similar preprocessing was done for the resting-state data, except for the smoothing. Tissue-type brain segmentation was run using FAST (Zhang et al., 2001), resulting in white-matter and ventricle masks for each participant. Functional images were aligned to the high-resolution anatomical volumes in each participant, initially using linear registration (FLIRT) and then optimized with Boundary-Based Registration (Greve & Fischl, 2009). Anatomical images were transformed to MNI space using FMRIB’s Nonlinear Image Registration Tool (FNIRT), and the resulting wrap fields were applied to the functional images in order to allow the projection of all participants onto a common brain template.

### Whole brain GLM analysis

Whole brain statistical maps were computed using a general linear model (GLM), separately for each of the three tasks. Voluntary and control blocks were used as the regressors, as well as the 6 standard and 18 extended motion parameters, as estimated by FSL MCFLIRT, in order to regress out motion artifacts. The regressors were convolved with the canonical double-gamma hemodynamic response function (HRF), attaining a model of the expected hemodynamic responses (Boynton et al., 1996). Multiple linear regression was performed for each run of each participant, obtaining estimates of the response amplitudes (beta values) in each voxel, for each of the conditions. The comparisons that were contrasted included the voluntary blocks (VF, AU and INST) vs. baseline, the control blocks vs. baseline, and the voluntary blocks vs. control blocks. Resulting contrast images were entered into a second-level analysis per participant, in order to combine the functional runs of each individual participant (fixed effects model). Next, group analysis was run using FMRIB Local Analysis of Mixed Effects (FLAME) (Smith et al., 2004), using the beta values of each participant as the dependent variables in a paired t-test in order to estimate inter-subject random effects. Resulting Z-statistic maps were corrected for multiple comparisons using cluster correction, determined by a voxel threshold of Z>2.3 and a corrected cluster significance threshold of p=0.05. Statistical maps were projected on inflated and flat cortical surfaces in MNI space, constructed using Freesurfer 5.3 (Dale et al., 1999; Fischl et al., 1999).

An additional GLM analysis was run in order to define individual participants’ ROIs, for further ROI analyses. It was identical to the GLM described above, only instead of modeling the entire blocks with a single predictor per block (for both voluntary and control blocks, in all three experiments), we created separate predictors for the initial 26 seconds of each block, and predictors for the remaining later part of each block. First level contrast images were entered into a second-level fixed-effect analysis, combining the separate functional runs of each participant. Individual ROI definition was based on the contrast of the initial 26 seconds of the voluntary blocks vs. baseline.

### ROI definition and time course extraction

Regions of interest (ROIs) were defined in order to carry out ROI analyses, examining the average responses and amplitude changes during the voluntary and control events, as well as the fALFF values during the verbal responses and resting state time courses, in the three different experiments. ROIs known to be involved in language processing, as well as in the three tasks employed here (verbal fluency, alternative uses and instances), were defined individually for each participant, including the left inferior frontal gyrus (IFG) and left middle frontal gyrus (MFG), separated to an anterior region (LH Pars Triangularis) and a posterior region (LH Pars Opercularis), the left precentral gyrus (LH PreCG), the paracingulate gyrus and anterior cingulate cortex (PCG and ACC), and the default mode network (DMN) regions (Schlosser et al., 1998; Costafreda et al., 2006; Beaty et al., 2014). All ROIs were defined individually for each participant, based on their activation maps in conjunction with atlas-based anatomical masks, in order to take into account the inter-subject variability in anatomical coordinates of functional regions. Critically, individual subjects’ activation maps were based on the contrast of the first 26 seconds of the voluntary blocks (VF/AU/INST) > baseline (breaks between blocks), while all further ROI-based analyses included only the events that occurred 30 seconds or later into the blocks, for both voluntary and control events. Thus, we avoid circularity in the definition and the analyses of the ROI data. Anatomical masks based on the Harvard-Oxford Cortical Structural Atlas (Desikan et al., 2006) were created for the LH Pars Triangularis, LH Pars Opercularis, LH precentral gyrus, LH insular cortex, paracingulate gyrus and anterior cingulate gyrus combined, and for the DMN regions, the precuneus and the posterior cingulate cortex (PCC) combined. An additional DMN region, the inferior parietal lobule (IPL), was defined based on the Julich histological atlas (Caspers et al., 2008). All ROIs, except for the two DMN regions, were defined as the voxels that showed increased activation in each subject’s contrast of the first 26 seconds of the voluntary blocks (VF/AU/INST) > baseline, thresholded at p<0.0001, in conjunction with the relevant anatomical masks. DMN ROIs were defined as the voxels that showed decreased activations in this contrast, using the same threshold. If no voxels survived the thresholding, it was gradually lowered until p<0.01, to obtain a minimal ROI of ∼30 voxels. If still no voxels survived, the ROI region was not defined for that participant. The LH anterior insula ROI was defined only for the verbal fluency task, and not for the alternative uses and instances task, as it did not survive the thresholding in most participants in these two tasks.

ROI time courses were extracted for each ROI in each participant, for the experimental runs as well as for the resting-state scans. Before the extraction of the ROI time courses, further motion correction procedures were done: in addition to regressing out the 6 standard and 18 extended motion parameters, estimated by FSL MCFLIRT, we also identified and excluded TRs with head movements using a scrubbing procedure (Power et al., 2012). Additionally, in the resting state data, non-neuronal contributions to the BOLD signal were removed by linear regression of motion parameters and ventricle and white-matter time courses from the unsmoothed data, for each participant (Fox et al., 2009; Hahamy et al., 2014). Following these steps, we averaged the BOLD signals across all voxels in each ROI, computing the time courses for each ROI in each participant.

### ROI event-related response analysis

In order to examine the event-related ROI responses, we first normalized the ROI BOLD time courses to percent signal change, relative to the mean BOLD amplitude across each entire run. The BOLD responses were locked to the onset of the voluntary and control verbal events, indicated by the button presses, excluding events that occurred in the first 30 seconds of each block, as explained in the *ROI definition and time course extraction* section above. Furthermore, only events that were separated by at least 12 “event-free” seconds from the previous event were analyzed, for both the voluntary and control conditions, thus allowing examination of the activity preceding these events, that is not contaminated by residual activity from previous events. Event-related responses were averaged across trials and experimental runs for each participant, separately for the voluntary and control conditions in the three experimental tasks, from −8 to +12 seconds relative to the button press. The BOLD responses were then averaged across all participants. The differential time courses of the two conditions were also calculated, defined as the within subject average voluntary response minus the control response, averaged across participants. Paired t-tests were conducted in each ROI and condition, comparing the BOLD amplitude at each time point to the baseline amplitude, defined as the average amplitude at times −6 and −4 relative to event onset.

In order to rule out the possibility that the gradual buildup preceding the voluntary events, but not the control events, is caused by an amplitude difference between these two conditions, we normalized the responses of individual participants to their peak value, using min-max normalization: x’= (x-min)/(max-min). This results in mapping the minimal value of the responses to 0 and the maximum value to 1, therefore obtaining an equal peak value of 1 in both the voluntary and control responses of each participant. After this normalization, we proceeded with the same pipeline as before, including averaging the responses across participants, calculating the differential time-courses, and conducting the statistical tests.

### Variance control simulation and analysis

A critical possible confound underlying the gradual buildup results is that this buildup was actually formed artificially by averaging across multiple “step-function” responses with time-jittered onsets, due to inaccuracies in the participants’ button press reports. In order to differentiate between a “true” single-trial-level gradual buildup and a time-jittered step-function alternative, we ran a variance simulation analysis, obtaining the expected variance time-courses of these two possible models, and then compared them to the real experimental data variance time-courses.

The simulation of each of the two models included 1000 iterations, with each iteration consisting of 30 simulated trials. For the jittered-time step function simulation, neural activity estimates for individual trial time-courses were constructed as box plot responses with 11 values, or “time-points”, in order to resemble the experimental responses we examined in the tasks, that consisted of 11 TRs, or 20 seconds (see *ROI event-related response analysis)*. These time points were marked between −8 and 12 seconds, solely for simple visual comparisons with the experimental data. These box plot responses began with an amplitude of 0±random noise, consisting of a random value between −0.35 to 0.35 (with 0.01 intervals), with a different random noise level for each time point and each trial. At a random time point, between −2 to 4 seconds relative to “event onset”, the amplitude abruptly increased to 1±random noise, remaining at this level until dropping back to ∼0 at time 6 seconds, in all trials. This resulted in obtaining an acro ss-trials averaged response that displayed a gradual buildup, starting at around time −2, and peaking at time 4 relative to event onset, matching the time dynamics in the real experimental voluntary event-related responses. Examples of 30 such simulated trials from one iteration are shown in the left panel of figure 7a, each trial marked in a different color, and the across-trial average in a thicker black line. The amplitude jumps from ∼0 to ∼1 are clearly seen at different time points. Next, we obtained the BOLD response estimates by convolving the neural estimate responses with the standard hemodynamic response function (Boynton et al 1996), and divided each response by its maximum value, to maintain the ∼0-1 amplitude scale. The BOLD estimates of the individual trials are shown in different colors in the middle panel of figure 7a, and the across trial average is shown in the black line.

In the single-trial gradual buildup simulation, the neural activity estimates had the same “time-duration” as in the jittered-time step function simulation, and also began with an amplitude of 0±random noise, consisting of a random value between −0.35 to 0.35 (with 0.01 intervals). Here, instead of a discrete step function occurring at a random time point, the responses demonstrated a triangular-shaped activity buildup, with a linear positive slope of 0.25, to which we added a random noise value of ±0.05 (with 0.01 intervals) in each simulated trial. The signal increase began at time −2, and reached a peak amplitude of 1±0.35 jittered noise, at time 4 relative to “event onset”. Thus, we obtain an across-trial average response that shows a slow accumulation from ∼-2 seconds, and peaks at time 4 relative to “event onset”, matching the experimental event-related responses in the 3 experiments. The left panel in figure 7b shows 30 example simulated trials from one iteration, each trial shown in a different color, with the across-trial average in black. BOLD responses estimates were obtained by convolving the responses with the standard HRF response, as was done in the step-function simulation, and these responses are shown in the middle panel of figure 7b. Comparing the left and middle panels of figure 7b with those in figure 7a, it is clear to see that the across-trial average in the step-function simulation and in the gradual-slope simulation are very similar, both displaying a gradual increase, yet one resulting from true single-trial gradual buildup, while the other results from a time-jitter in the abrupt step-function.

The variance time-courses for the two models were obtained by calculating the across-trial variance in each iteration (across the 30 trials), for each time point, and then averaging the variance values across the 1000 iterations. In order to evaluate changes in the variance along the time-course, the variance value at each time point was compared to the baseline variance, defined as the average variance at times −6 and −4 before “event onset”, similar to the comparisons done in the mean amplitude event-related responses (see *ROI event-related analysis).* P-values for each time point were obtained from the distribution of 1000 variance difference values between the variance and the mean baseline variance (one difference value from each iteration). The p-values were defined as 2*minimum(P_val, 1-P_val), with P_val defined as the proportion of variance difference values smaller than 0.

The variance time-courses of the actual experimental data were calculated for each of the three experiments, separately for the voluntary and control conditions, in each ROI, as well as for the pupil. The across-trial variance time course of each participant was calculated from the same trials as explained in *ROI event-related response analysis*, and then averaged across participants. A one-tailed paired t-test was calculated for each time point, examining whether the variance at each time point was higher than the baseline variance (average variance at times −6 and −4 seconds before event onset).

### fALFF analysis and simulation

In order to quantify and compare the time-frequency dynamics during the resting state, to those of the voluntary and control responses, we calculated the fractional amplitude of low frequency fluctuations (fALFF) (Zou et al., 2008; Zuo et al., 2010) in these conditions for each participant. The fALFF values of the voluntary and control responses were calculated from the average event-related response of each participant, for all the ROIs defined, separately for the 3 tasks (VF, AU, and INST). These individual averaged event-related responses were obtained as described in the *ROI event-related response analysis* section, following the preprocessing steps described in *fMRI preprocessing* and *ROI definition and time-course extraction,* and percent signal change normalization. These mean responses, for each participant, each ROI, and each condition, were transformed to the power-frequency domain using the Fourier frequency transform. Next, the sum of the square root of the power amplitudes across the 0.01-0.1 Hz frequency range was divided by the sum of the square root of the power across the entire frequency range measured (0-0.25 Hz). This resulted in an fALFF value for each subject and each ROI, for the mean VF, AU, and INST responses, and their matching control responses. Similarly, we obtained the fALFF values of the resting state time course of each subject, for the two resting state scans that were performed (prior to the VF experiment, and prior to the AU and INST experiments, as explained in *The experimental tasks and design* section). Resting state data first underwent preprocessing, including regressing out the motion parameters, white matter and ventricle time-courses, as described in *fMRI preprocessing* and *ROI definition and time course extraction*. Next, the resting state time-courses were transformed to the power-frequency domain, by Fourier Transform, and the square root of the power was calculated for each frequency. Then, the sum of the square root amplitudes across 0.01-0.1 Hz was divided by the sum across the entire frequency range, resulting in an fALFF value for each participant and each ROI.

Finally, the resting state fALFF values were correlated with the voluntary and the control fALFF values, separately for the VF, AU and INST tasks, using Spearman correlation. All correlation p-values were derived from a non-parametric permutations test, with 10,000 random subject-wise permutations, followed by FDR correction at α=0.05, to correct for multiple comparisons across the different ROIs.

fALFF calculations are usually commonly for depicting resting state data (Zou et al., 2008; Zuo et al., 2010). However, here we opted to use this measure for examining average event-related responses as well. In order to validate this usage, we run a simulation, in which we generated a series of 17 “resting-state” and “event-related” responses, with common slopes, or wave-shapes, and examined their fALFF measures.

“Event-related responses” were created in a similar manner to the responses generated in the variance simulation, as described in *Variance control simulation and analysis.* Neural activity estimates were constructed as vectors with 30 elements, with all values ranging between 0 and 1. All responses began with an amplitude of 0, and then started linearly increasing in a triangular-shaped buildup with a different positive slope in each simulated response, ranging between 0.05 to 0.5 (with a total of 17 different slopes). After reaching a peak amplitude of 1, the next value in the series dropped back to 0. Examples for two simulated “event-related responses”, with a gradual and a steep slope, are shown in figure S9a. In order to obtain the BOLD response estimates, the responses were then convolved with the standard HRF response (Boynton et al 1996), also shown in figure S9a.

Simulated “resting state time courses” were generated by concatenating individual “event-related responses” with a constant slope, obtaining a vector with a total length of 360 values, as in the real experimental resting state data. Intervals of 0-amplitude elements with varying lengths, between 0 to 20 elements, separated between two consecutive waves. Additionally, random overlaps between two consecutive waves were also inserted, with random overlap inserts of between −10 to 0 steps, or elements, thus allowing sporadic summation of two successive simulated waves. BOLD response estimates were obtained by convolving these time courses with the standard HRF. Figure S9a shows two examples of the neural estimates and BOLD estimates of simulated resting state time courses with two different slopes. In the same manner as for the simulated “event-related responses”, we generated a total of 17 simulated “resting state time-courses” with varying buildup slopes, ranging between 0.05-0.5. Thus, for every slope value, we had generated one simulated “resting state time course”, and one simulated “event-related response”.

Next, we calculated the fALFF value for each simulated resting state time course and event related time course, in the same manner that was done for the experimental data (as described above). In order to inspect the relationship between the slopes from which the time courses were generated, and the fALFF values, we correlated between the two parameters, both for the resting-state and the event-related simulated data separately, using Spearman correlation. Additionally, we checked for a correlation between the fALFF values of the resting state and event related simulated time courses that were generated with a common slope, using Spearman correlation. All correlation plots are shown in figure S9b.

## Acknowledgments

The authors wish to thank Dr. Edna Furman-Haran, Fanny Attar and Eiska Tegareh at the Weizmann MRI center for their assistance. This study was funded by the Joy Ventures grant and CIFAR Tanenboum Fellowship to R.M.

