## Supplementary Figures for "Resting state fluctuations underlie free and creative verbal behaviors in the human brain"

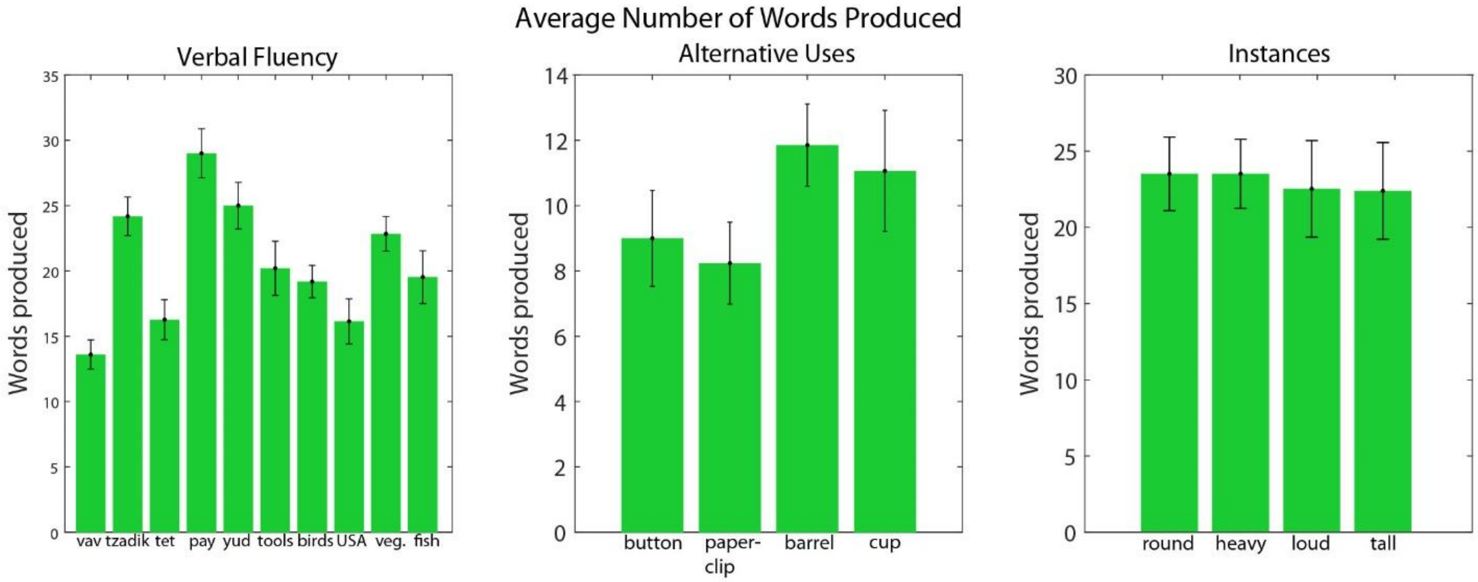


**Figure S1**. Mean number of words produced in the three experimental tasks, averaged across participants. Error bars represent the standard error.

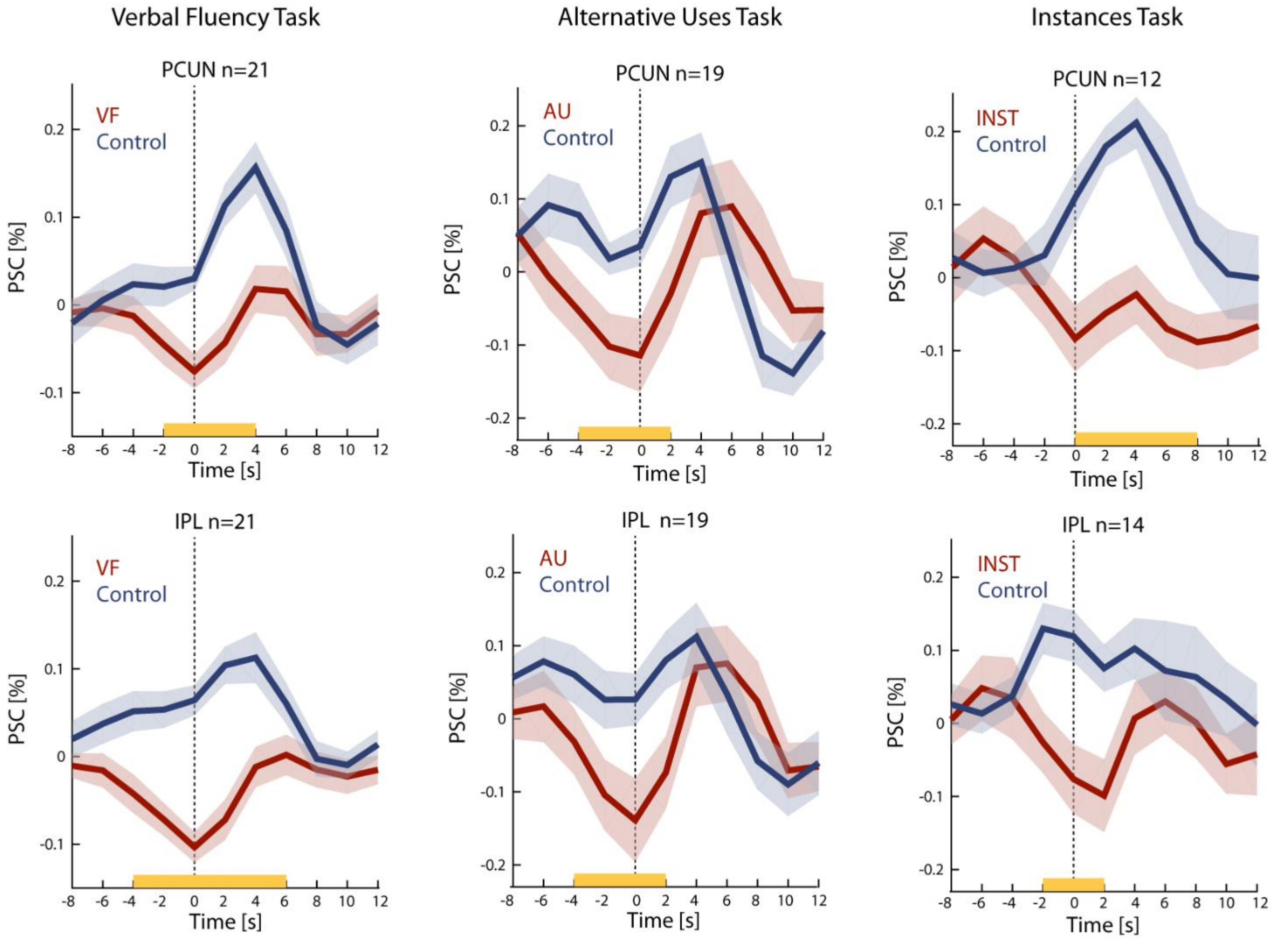


**Figure S2**. DMN ROIs group mean time courses during the voluntary and control conditions in the VF, AU and INST tasks. Vertical dashed lines indicate the report of a verbal generation event (button press). For information regarding individual ROI definition, see *ROI definition and time-course extraction* in *Methods*. Transparent borders indicate the mean $\pm$SE. Yellow lines indicate a significant difference between the two conditions (two-tailed paired t-test, $p<0.05$, cluster correction).

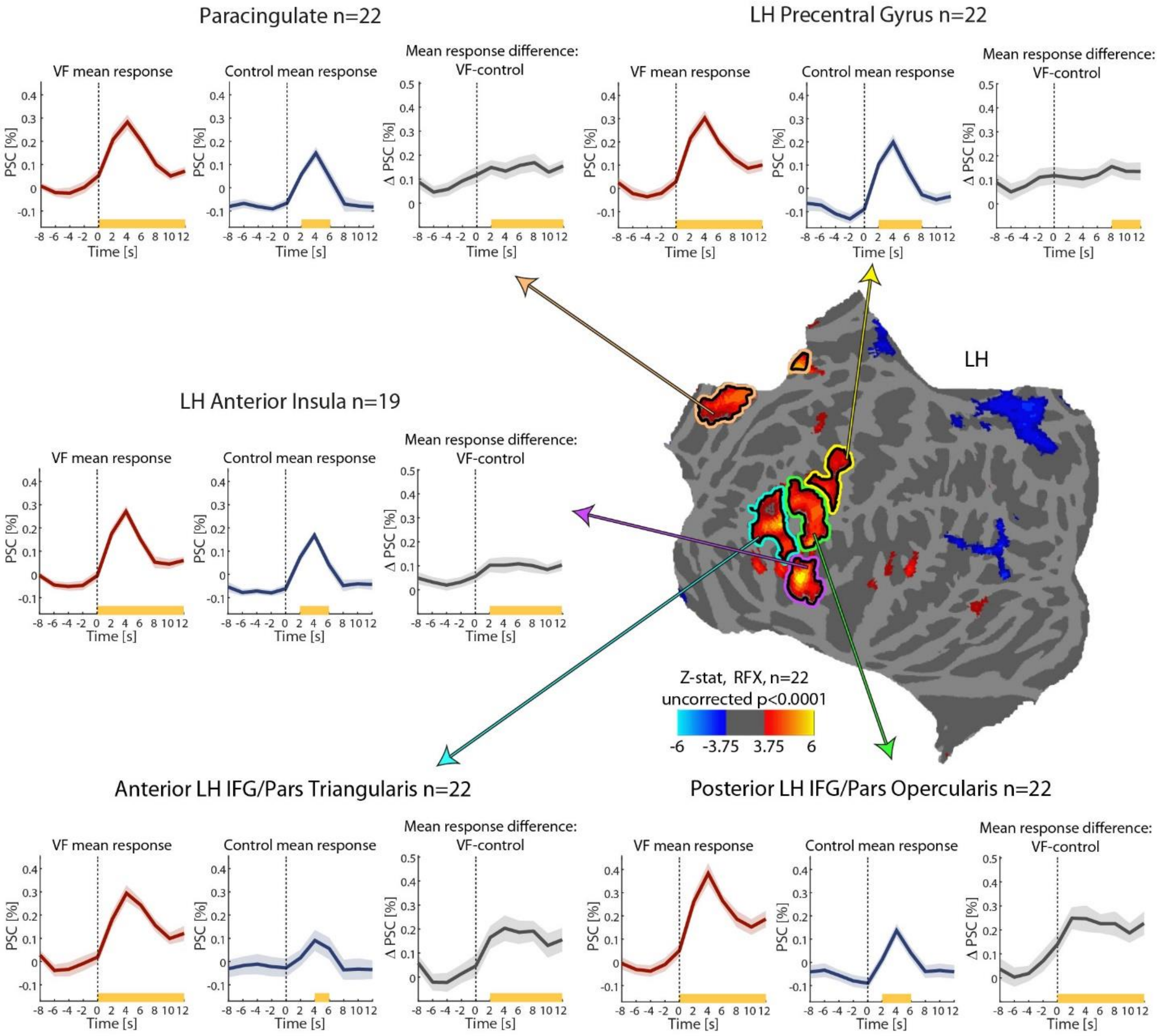


**Figure S3**. Group averaged BOLD responses during verbal fluency and control word generation events in 5 ROIs. Flat left hemisphere displays the group contrast of the first 30 seconds of the VF blocks >baseline, thresholded at $p<0.0001$ uncorrected, shown for ROI definition illustration purposes. Colored outlines denote ROIs defined by the group maps, solely for visualization purposes, while actual ROIs were defined individually for each participant, based on this contrast at the single subject level in conjunction with anatomical masks (see *ROI definition and time-course extraction* in the *Online Methods* for details). The anterior LH IFG is outlined in cyan, the posterior LH IFG in green, the LH anterior insula in pink, the LH precentral gyrus in yellow and the paracingulate cortex in beige-orange. Mean percent signal changes in the BOLD signal, averaged across participants, are presented in red for the verbal fluency events, blue for the control events, and gray for the mean response difference, defined as the verbal fluency event time-course minus the control time-course. Transparent borders indicate the mean ±SE. Dashed vertical lines indicate the time of the button press, reporting a verbal generation event (voluntary and control). Yellow lines indicate a significant increase in response amplitude above baseline, defined as the average amplitude across times -6 and -4 seconds before event onsets (one-tailed paired t-test, $p<0.05$).

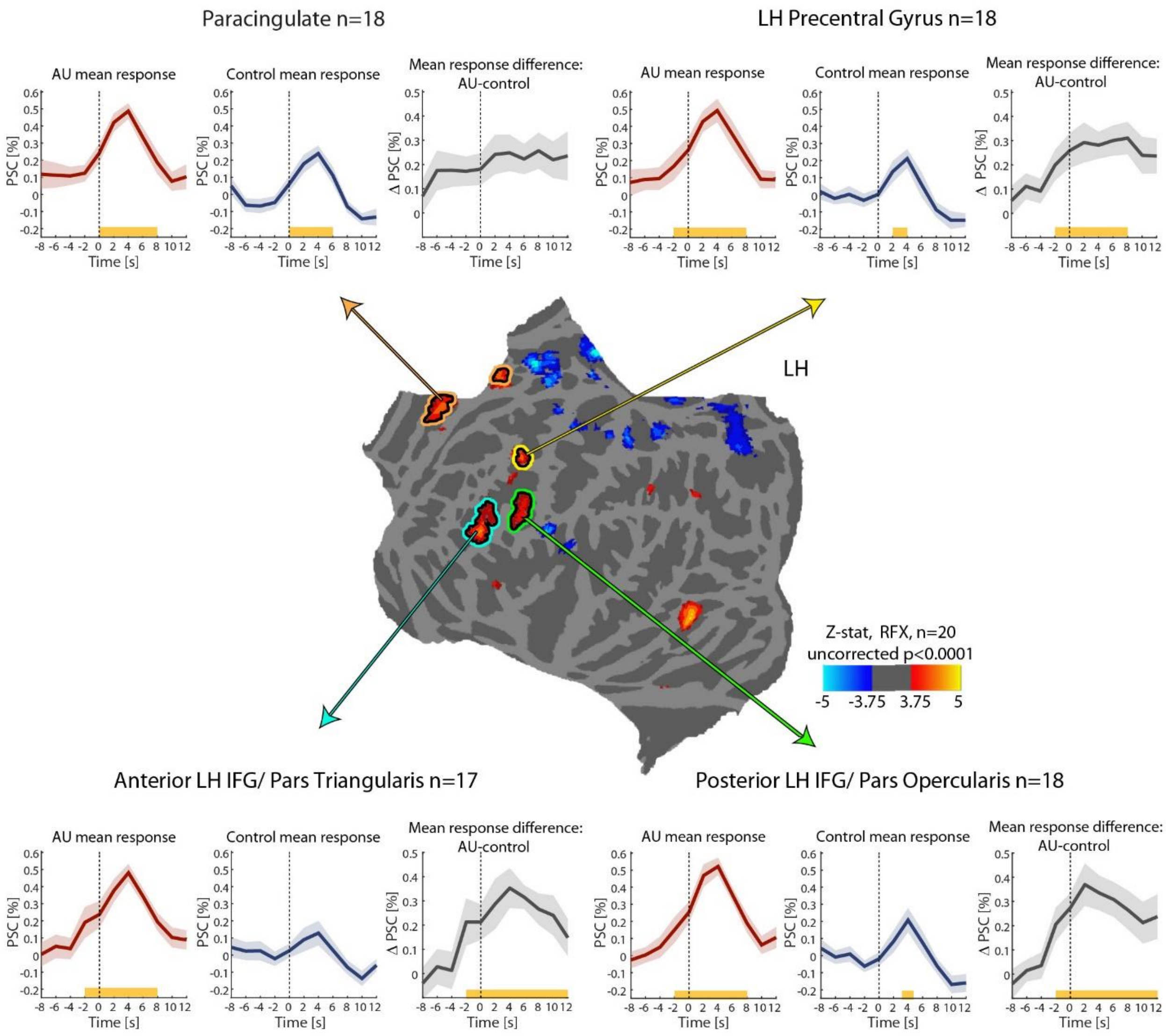


**Figure S4**. Group averaged BOLD responses during alternative uses and control events in 4 ROIs. Flat left hemisphere displays the group contrast of the first 30 seconds of the AU blocks >baseline, thresholded at $p<0.0001$ uncorrected, shown for ROI definition illustration purposes. Colored outlines denote ROIs defined by the group maps, solely for visualization purposes, while actual ROIs were defined individually for each participant, based on this contrast at the single subject level in conjunction with anatomical masks (see *ROI definition and time-course extraction* in *Online Methods* for details). The anterior LH IFG is outlined in cyan, the posterior LH IFG in green, the LH precentral gyrus in yellow and the paracingulate cortex in beige-orange. Mean percent signal changes in the BOLD signal, averaged across participants, are presented in red for the AU events, blue for the control events, and gray for the mean response difference, defined as the AU event time-course minus the control time-course. Transparent borders indicate the mean ±SE. Dashed vertical lines indicate the time of the button press, reporting a verbal generation event (voluntary and control). Yellow lines indicate a significant increase in response amplitude above baseline, defined as the average amplitude across times -6 and -4 seconds before event onsets (one-tailed paired t-test, $p<0.05$).

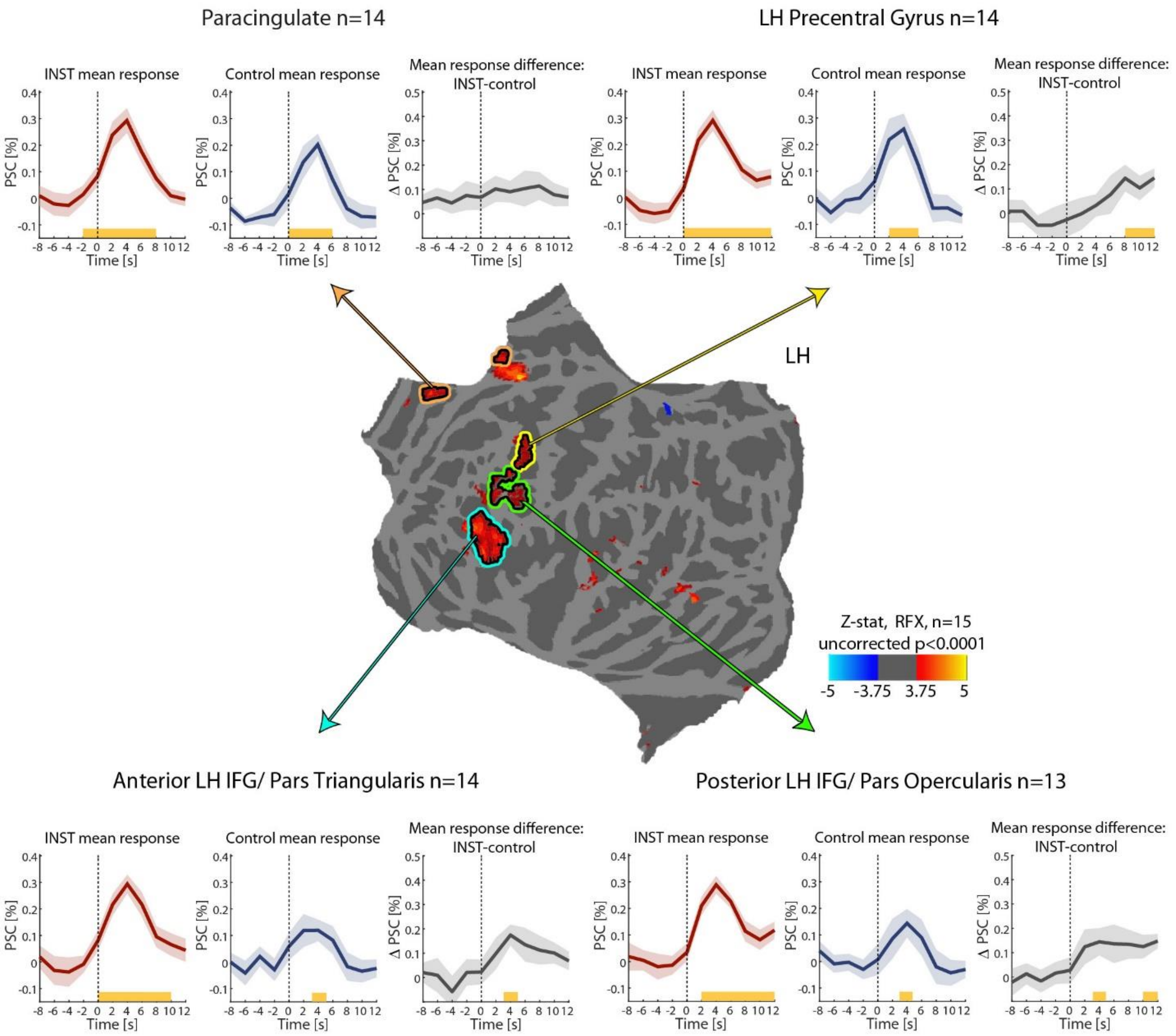


**Figure S5**. Group averaged BOLD responses during instances and control word generation events in 4 ROIs. Flat left hemisphere displays the group contrast of the first 30 seconds of the INST blocks >baseline, thresholded at p<0.0001 uncorrected, shown for ROI definition illustration purposes. Colored outlines denote ROIs defined by the group maps, solely for visualization purposes, while actual ROIs were defined individually for each *participant, based* on this contrast at the single subject level in conjunction with anatomical masks (see *ROI definition and time-course extraction* in *Online Methods* for details). The anterior LH IFG is outlined in cyan, the posterior LH IFG in green, the LH precentral gyrus in yellow and the paracingulate cortex in orange-beige. Mean percent signal changes in the BOLD signal, averaged across participants, are presented in red for the INST events, blue for the control events, and gray for the mean response difference, defined as the INST event time-course minus the control time-course. Transparent borders indicate the mean ±SE. Dashed vertical lines indicate the time of the button press, reporting a verbal generation event (voluntary and control). Yellow lines indicate a significant increase in response amplitude above baseline, defined as the average amplitude across times -6 and -4 seconds before event onsets (one-tailed paired t-test, p<0.05).

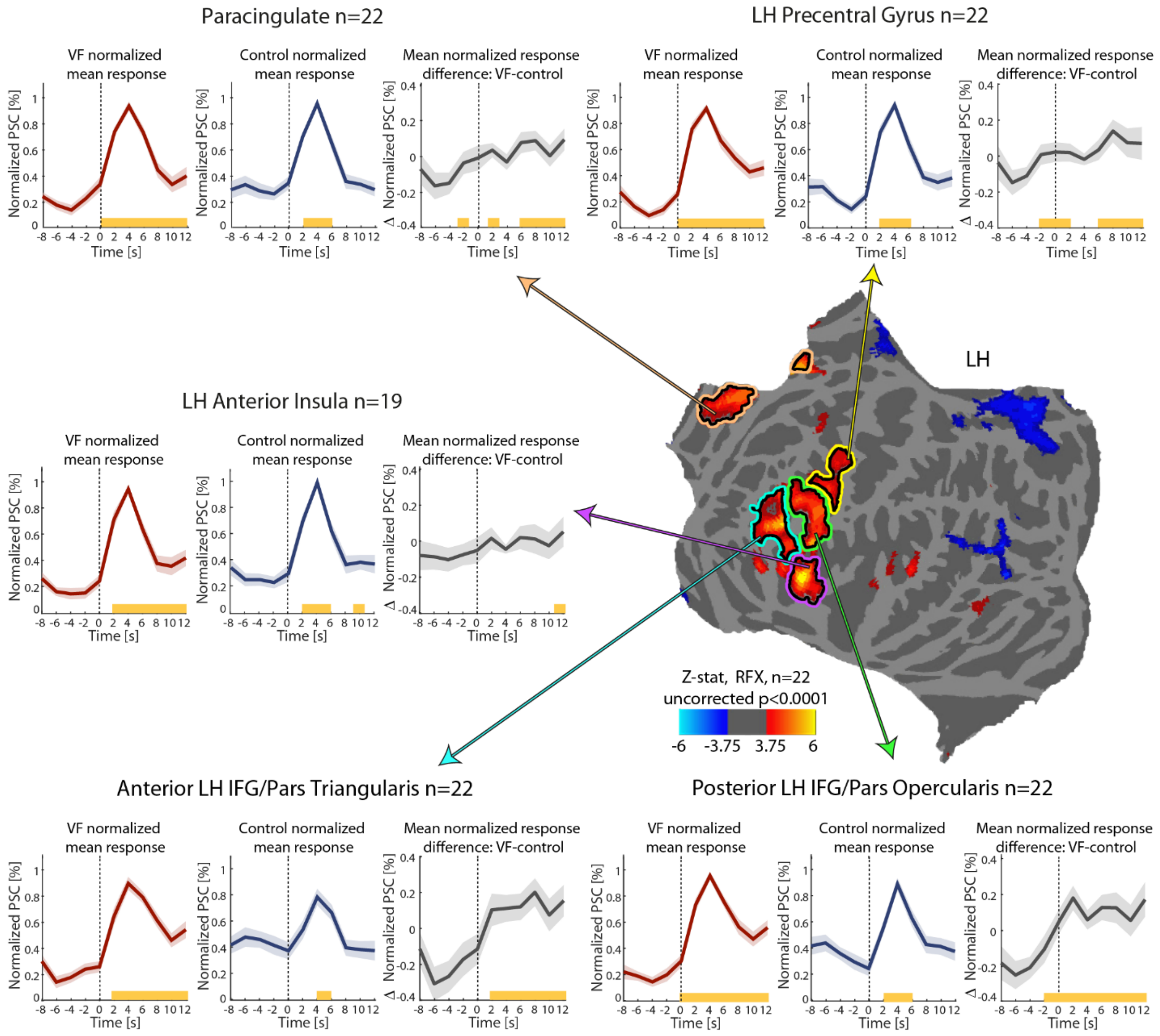


**Figure S6**. Min-max normalized average BOLD responses during verbal fluency and control word generation events in 5 ROIs. Individual participants' responses for the VF and control conditions were normalized separately using min-max normalization ((x-min)/(max-min), see *ROI event-related response analysis* in *Online Methods*), so that the new minimal value in each response time-course is 0, and the new maximal value (the peak amplitude) is 1. The normalized responses were then averaged across participants. Additional details are identical to those specified in figure S3.

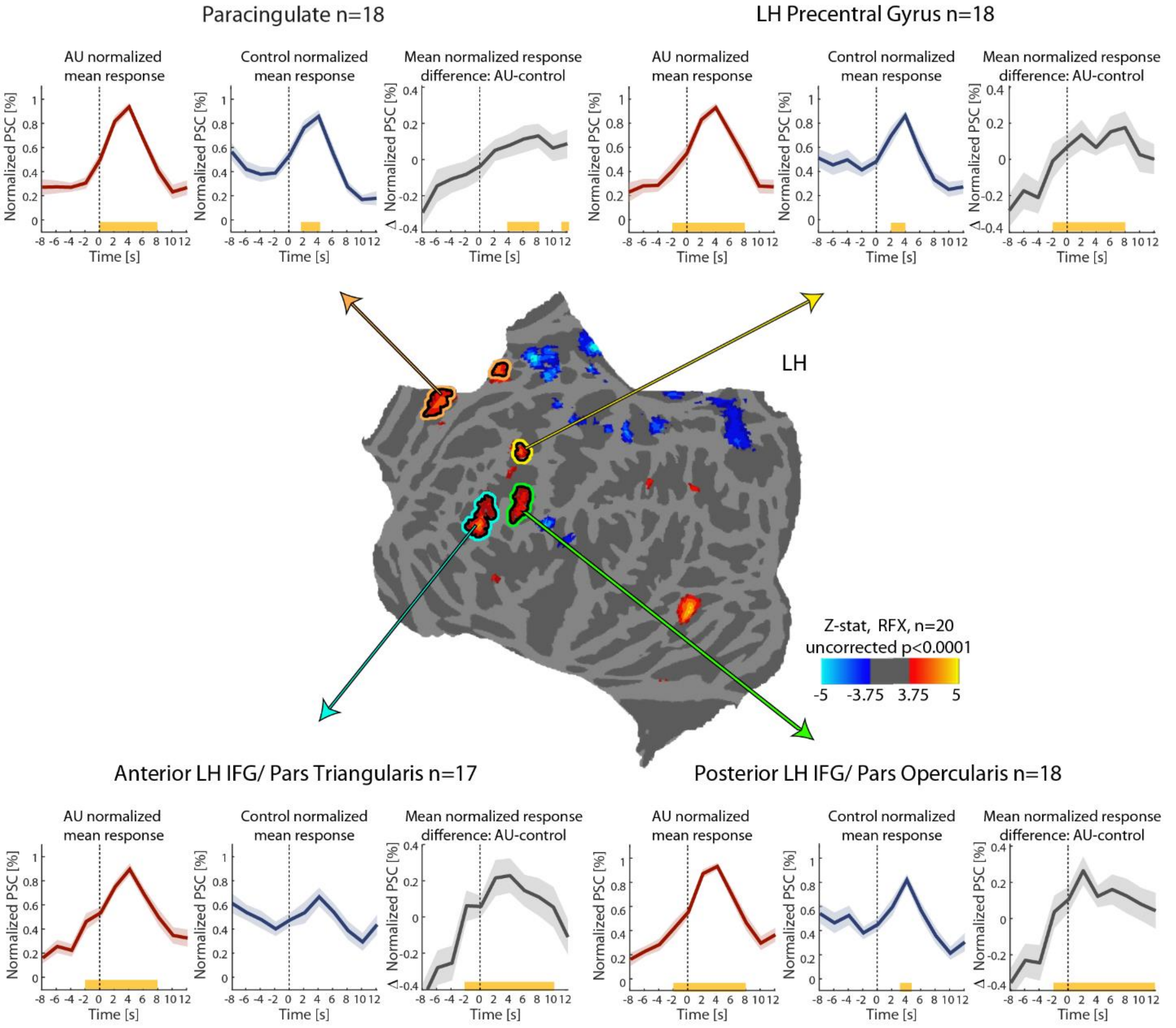


**Figure S7**. Min-max normalized average BOLD responses during alternative uses and control word generation events in 4 ROIs. Individual participants' responses for the AU and control conditions were normalized separately using min-max normalization ((x-min)/(max-min), see *ROI event-related response analysis* in *Online Methods*), so that the new minimal value in each response time-course is 0, and the new maximal value (the peak amplitude) is 1. The normalized responses were then averaged across participants. Additional details are identical to those specified in figure S4.

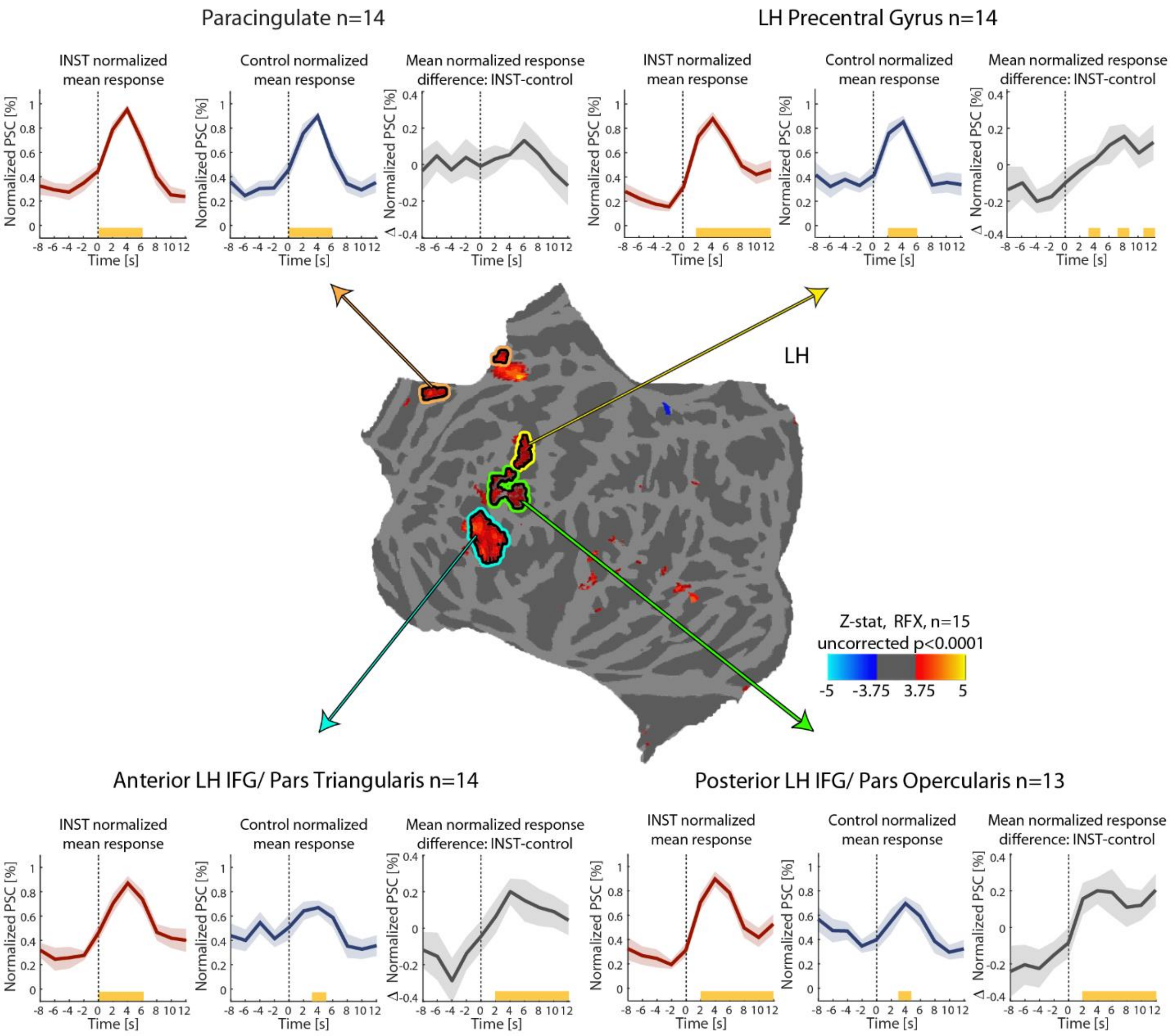


**Figure S8**. Min-max normalized average BOLD responses during instances and control word generation events in 4 ROIs. Individual participants' responses for the INST and control conditions were normalized separately using min-max normalization ((x-min)/(max-min), see *ROI event-related response analysis* in *Online Methods*), so that the new minimal value in each response time-course is 0, and the new maximal value (the peak amplitude) is 1. The normalized responses were then averaged across participants. Additional details are identical to those specified in figure S5.

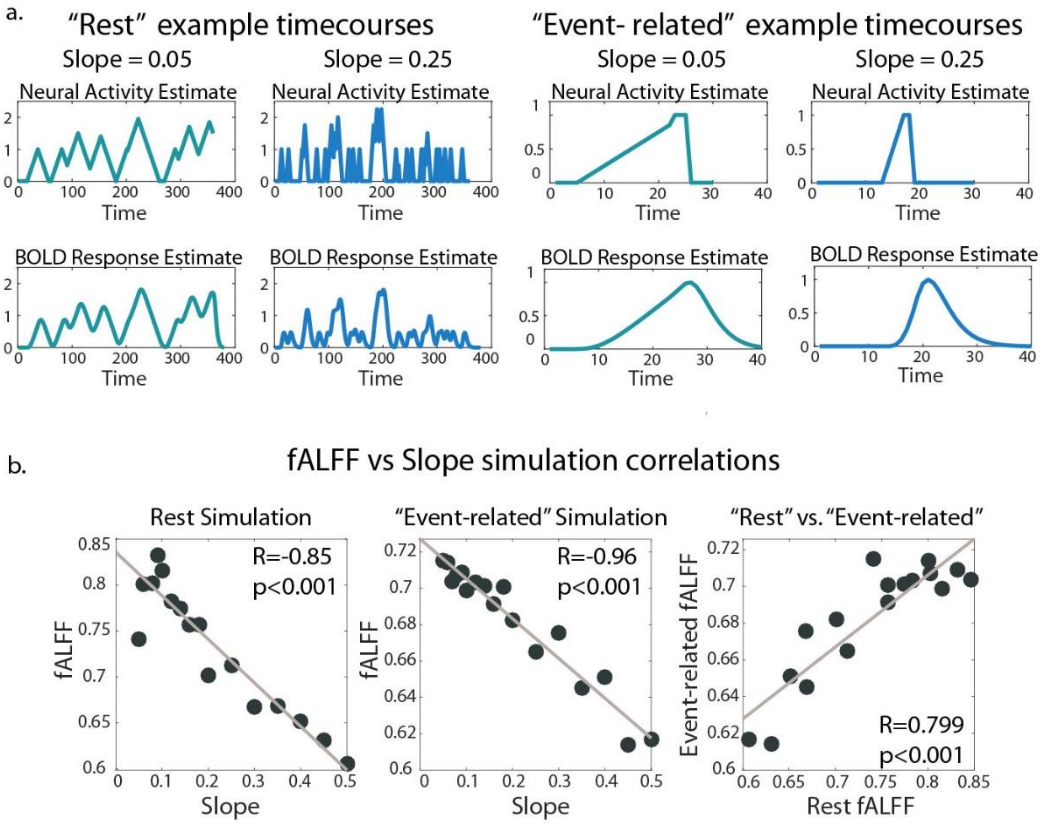


**Figure S9**. A. Examples of simulated neural and BOLD responses with two different slopes, a "slow" slope (0.05) and a "fast" slope (0.25), for "resting state" and "event-related" signals. B. Significant correlations between the fALFF values and the slopes of simulated resting state time series (left panel), between the fALFF and the slopes of event-related simulated responses (middle panel), and between the fALFF values of the "event-related responses" and "resting state time-courses" with common slopes (left panel). For additional details, see *fALFF analysis and simulation* in *Online Methods*.

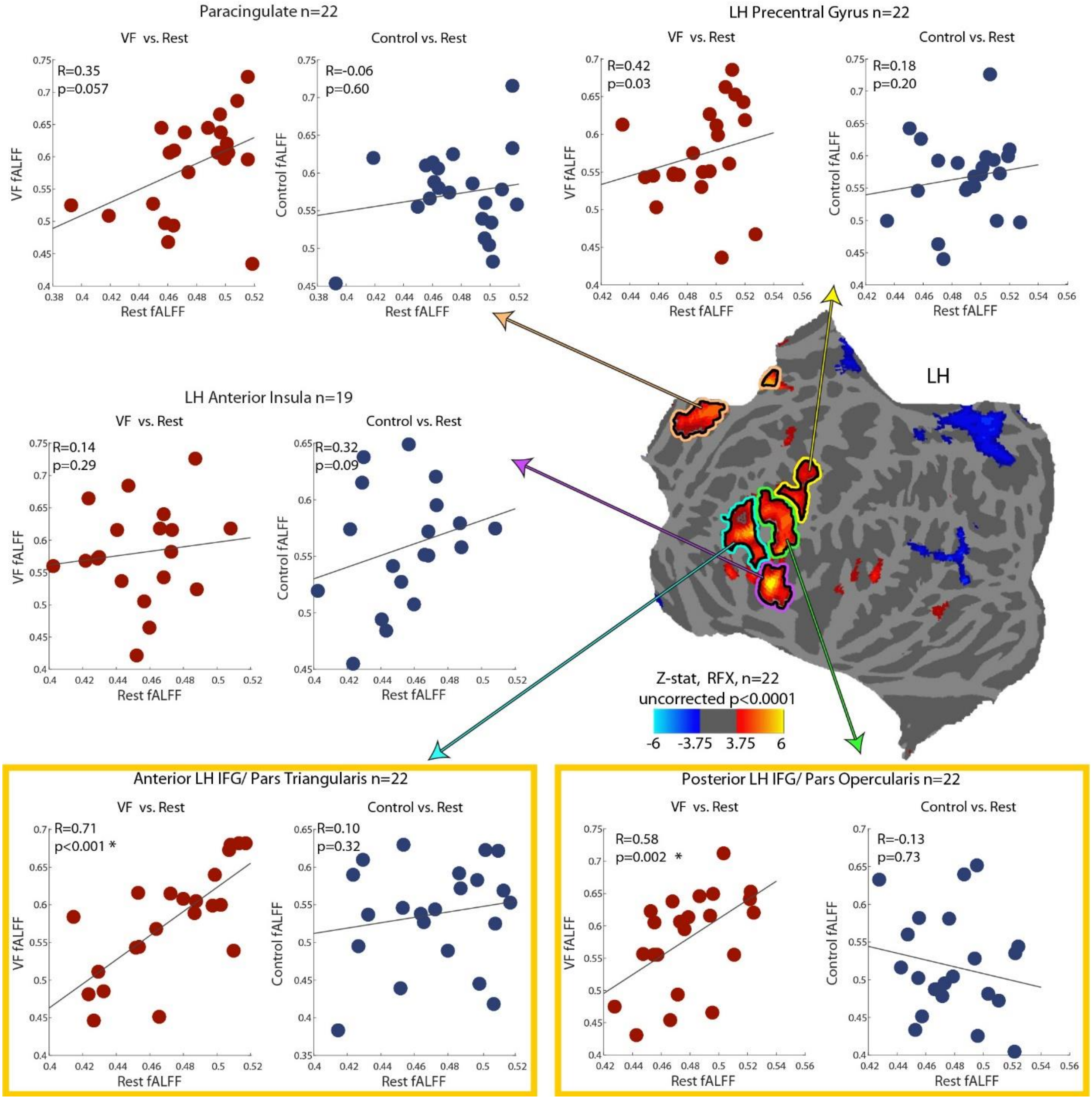


**Figure S10**. VF task fALFF correlation plots, depicting the correlations between the fALFF values of VF average responses vs resting state time courses, and control average responses vs resting state, in 5 ROIs. The same cortical map shown in figures S3 and S6 is presented here, for ROI illustration purposes. Left panels (red markers) depict the correlation between the VF and rest fALFF values, while the right panels (blue dots) depict correlations between control and rest. Every dot in the scatter plots represents one participant. Spearman's R correlation coefficients are presented, together with their p-values, derived from a subject-wise permutation test (10,000 permutations). Significant correlations are marked with an asterisk (p<0.05, FDR correction). Yellow frames mark a significant correlation difference between the VF and control correlations with rest, p<0.05, dependent correlation percentile bootstrapping test (Wilcox, 2016).

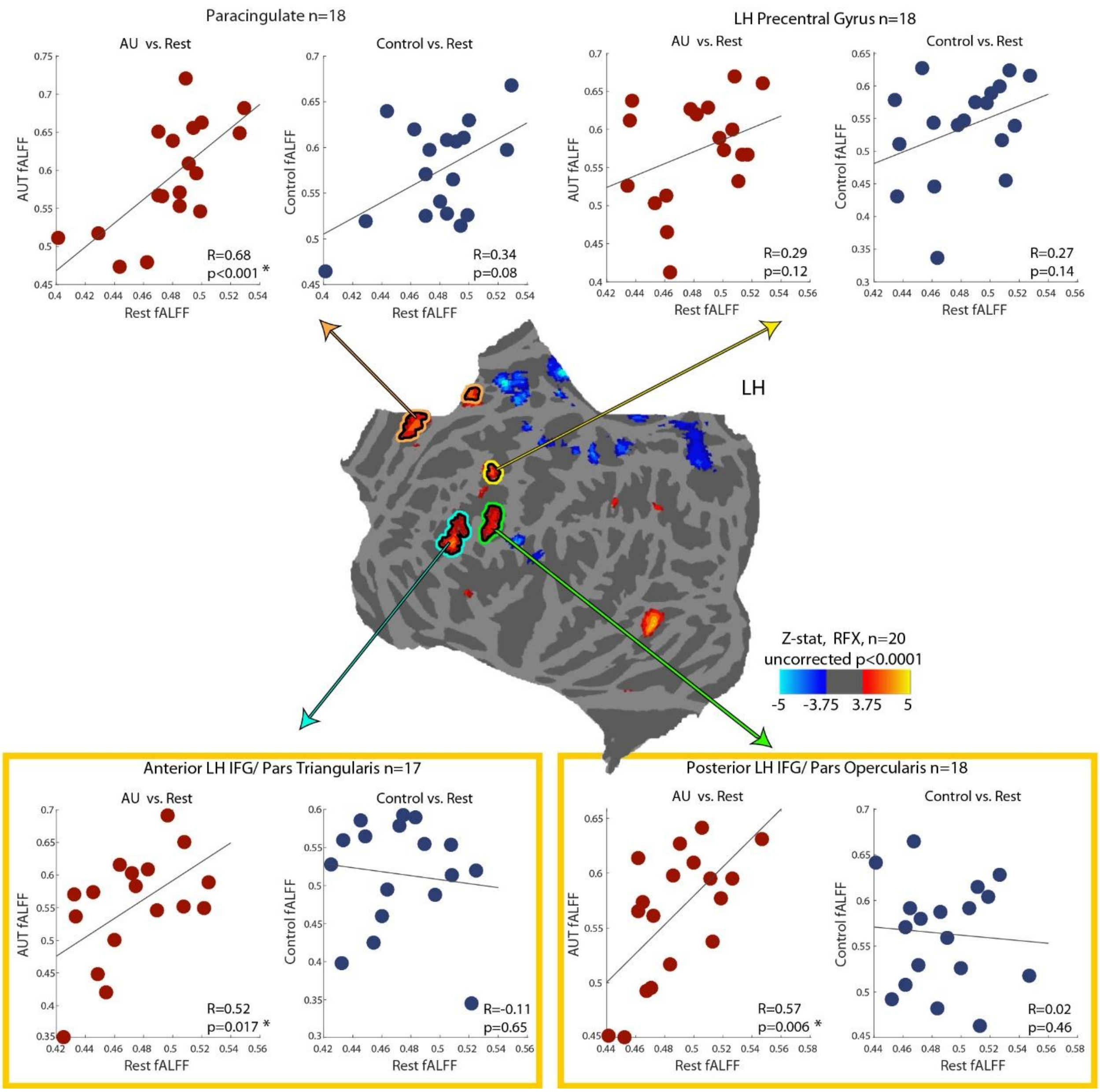


**Figure S11**. AU task fALFF correlation plots, depicting the correlations between the fALFF values of AU average responses vs resting state time courses, and control average responses vs resting state, in 4 ROIs. The same cortical map shown in figures S4 and S7 is presented here, for ROI illustration purposes. Left panels (red markers) depict the correlation between the AU and rest fALFF values, while the right panels (blue dots) depict correlations between control and rest. Every dot in the scatter plots represents one participant. Spearman's R correlation coefficients are presented, together with their p-values, derived from a subject-wise permutation test (10,000 permutations). Significant correlations are marked with an asterisk (p<0.05, FDR correction). Yellow frames mark a significant correlation difference between the AU and control correlations with rest, p<0.05, dependent correlation percentile bootstrapping test (Wilcox, 2016).

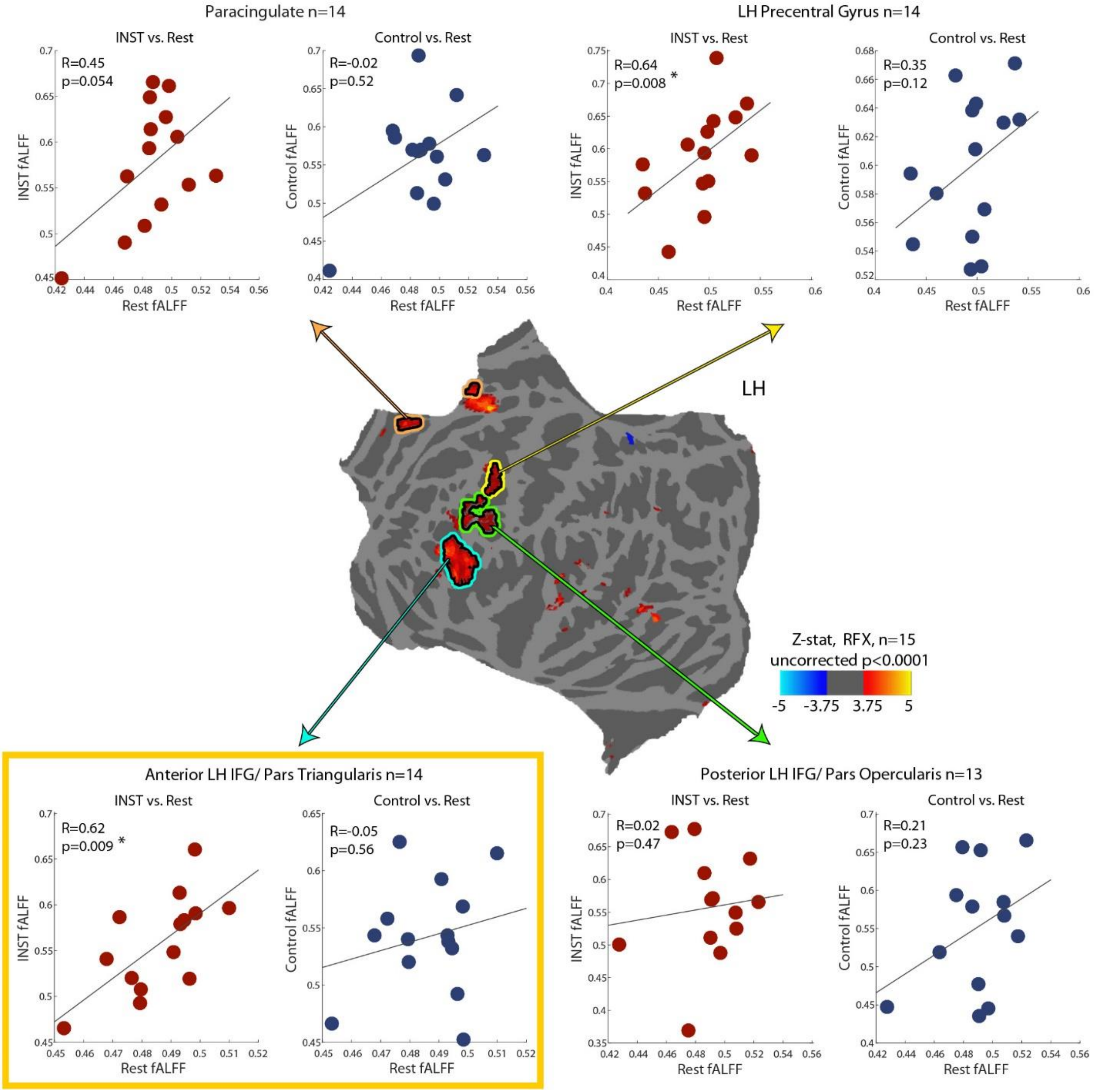


**Figure S12**. INST task fALFF correlation plots, depicting the correlation between the fALFF values of INST average responses vs resting state time courses, and control average responses vs resting state, in 4 ROIs. The same cortical map shown in figures S5 and S8 is presented here, for ROI illustration purposes. Left panels (red markers) depict the correlation between the INST and rest fALFF values, while the right panels (blue dots) depict correlations between control and rest. Every dot in the scatter plots represents one participant. Spearman's R correlation coefficients are presented, together with their p-values, derived from a subject-wise permutation test (10,000 permutations). Significant correlations are marked with an asterisk (p<0.05, FDR correction). Yellow frames mark a significant correlation difference between the INST and control correlation with rest, p<0.05, dependent correlation percentile bootstrapping test (Wilcox, 2016).
